# Cortical Contributions to Medial Frontal β-Bursts during Executive Control

**DOI:** 10.1101/2022.10.04.510901

**Authors:** Steven P. Errington, Jacob A. Westerberg, Geoffrey F. Woodman, Jeffrey D. Schall

**Affiliations:** Department of Psychology, Vanderbilt Vision Research Center, Center for Integrative & Cognitive Neuroscience, Vanderbilt Brain Institute, Vanderbilt University, Nashville, Tennessee 37240, USA; Centre for Vision Research, Vision: Science to Applications Program, Department of Biology, York University, Toronto, ON M3J 1P3, Canada

## Abstract

EEG β-bursts observed over the medial frontal cortex are claimed to mediate response inhibition despite their infrequent occurrence. The weak association with stopping behavior is supposed to be a by-product of the low signal-to-noise ratio of EEG recordings. We tested the premise that β-bursts are more common within the cerebral cortex and more directly associated with response inhibition. We sampled simultaneously EEG and intracortical local field potentials (LFP) within the medial frontal cortex (MFC) of two macaque monkeys performing a response inhibition task. Intracortical β-bursts were just as infrequent as those in EEG and did not parallel the likelihood of canceling a planned response. Cortical β-bursts were more prevalent in upper layers but were not synchronized across a cortical column or with EEG β-bursts. These findings contradict claims for a causal contribution of β-bursts during response inhibition, provide important constraints for biophysical and cortical circuit models, and invite further considerations of β-burst function in cognitive control.

## INTRODUCTION

Cognitive control describes a set of processes, associated with the frontal cortex and related brain networks, that are important when automatic or learned actions are detrimental to achieving a goal (Logan and Cowan, 1984). Inappropriate actions must be stopped before execution to avoid interference with task goals. This feature of cognitive control is referred to as response inhibition. Recent work in humans has proposed that β-bursts in the frontal cortex enable response inhibition (Enz et al., 2021; Hannah et al., 2020; Jana et al., 2020; Wessel, 2020). However, we recently showed that EEG β-bursts occurred too infrequently, did not parallel stopping behavior, and multiplexed other cognitive functions (Errington et al., 2020).

Several explanations have been offered for these divergent observations. First, the rarity of β-bursts and associated lack of causal efficacy for stopping has been attributed to the low signal-to-noise ratio of scalp EEG (Hannah et al., 2020; Jana et al., 2020; Wessel, 2020). Second, although conclusions drawn from human studies have emphasized evidence that medial frontal areas contribute to the interruption of response preparation (Diesburg and Wessel, 2021; Rae et al., 2014; Sharp et al., 2010), the conclusions are refuted by neurophysiological evidence obtained in the medial frontal cortex of macaques (Scangos and Stuphorn, 2010; Stuphorn et al., 2010). Finally, although β-bursts have been observed in EEG contacts directly over areas in the medial frontal cortex, EEG signals reflect complex contributions from multiple areas (Naess et al., 2021).

Although the relationship between intracortical LFP and EEG signals is uncertain, we know that EEG signals measured over the scalp are generated by microcircuits within the cortex (e.g., Herrera et al. 2020; 2022) such that associations between the activity of particular neurons and EEG signatures can be observed (e.g., Sajad et al. 2019, 2022). Biophysical modeling has suggested that cortical β-bursts can be produced via a transient dipole reversal caused by the interaction between a prolonged excitatory synaptic drive in the basal dendrites of pyramidal neurons that interacts with a transient pulse of excitatory drive in the apical dendrites (Sherman et al., 2016; Shin et al., 2017). However, it remains to be tested whether such layer-specific activity is observed in the medial frontal cortex, and how this contributes to EEG signals recorded from the overlying cranial surface.

We addressed these issues by simultaneously recording LFP across all cortical layers of the Supplementary Eye Field and from EEG electrodes placed on top of the skull in monkeys performing a saccade countermanding task. This task has a strong theoretical framework through which neural activity related to reactive or proactive control can be distinguished (Boucher et al., 2007a; Lo et al., 2009; Logan and Cowan, 1984; Logan et al., 2015; Wiecki and Frank, 2013).

Despite the increase in signal-to-noise ratio, we observed intracortical β-bursts as infrequently as those in the EEG. LFP β-bursts were more common during stopping but lacked causal efficacy. LFP β-bursts were also prevalent after execution of the response and during anticipation of reward. LFP β-bursts were most prominent in the upper cortical layers but were not synchronized across the cortical column. Although SEF exhibits the functional, biophysical, and anatomical properties required to contribute to EEG signals (Sajad et al. 2019, 2022; Herrera et al. 2020), we found no evidence that EEG β-bursts could be caused by LFP β-bursts. These findings challenge a prevailing hypothesis and constrain models of the functional microcircuitry of β-burst generation during cognitive control.

## RESULTS

### Countermanding performance and neural sampling

We acquired 33,816 trials across 29 sessions from two macaques (Eu: 11,583 trials; X: 22,233 trials) performing the saccade countermanding task (**Fig. 1A**). Both monkeys exhibited typical sensitivity to the stop-signal (**Supplementary Fig. 1A**). First, response latencies on non-canceled (error) trials were faster than those on no-stop trials. Secondly, the probability of failing to inhibit a saccade was greater at longer stop-signal delays. These two observations validated the assumptions of the independent race model (Logan & Cowan, 1984), allowing us to estimate the stop-signal reaction time (SSRT). Monkeys had indistinguishable mean SSRT (one-way independent measure ANOVA: F (1, 27) = 0.108, p = 0.745) and variance of SSRT (one-way independent measures ANOVA: F (1, 27) = 0.819, p = 0.819). However, trigger failures were significantly more common for monkey Eu relative to monkey X (one-way independent measures ANOVA: F (1, 27) = 18.458, p < 0.001, BF_10_ = 114.778).

**Fig. 1.**
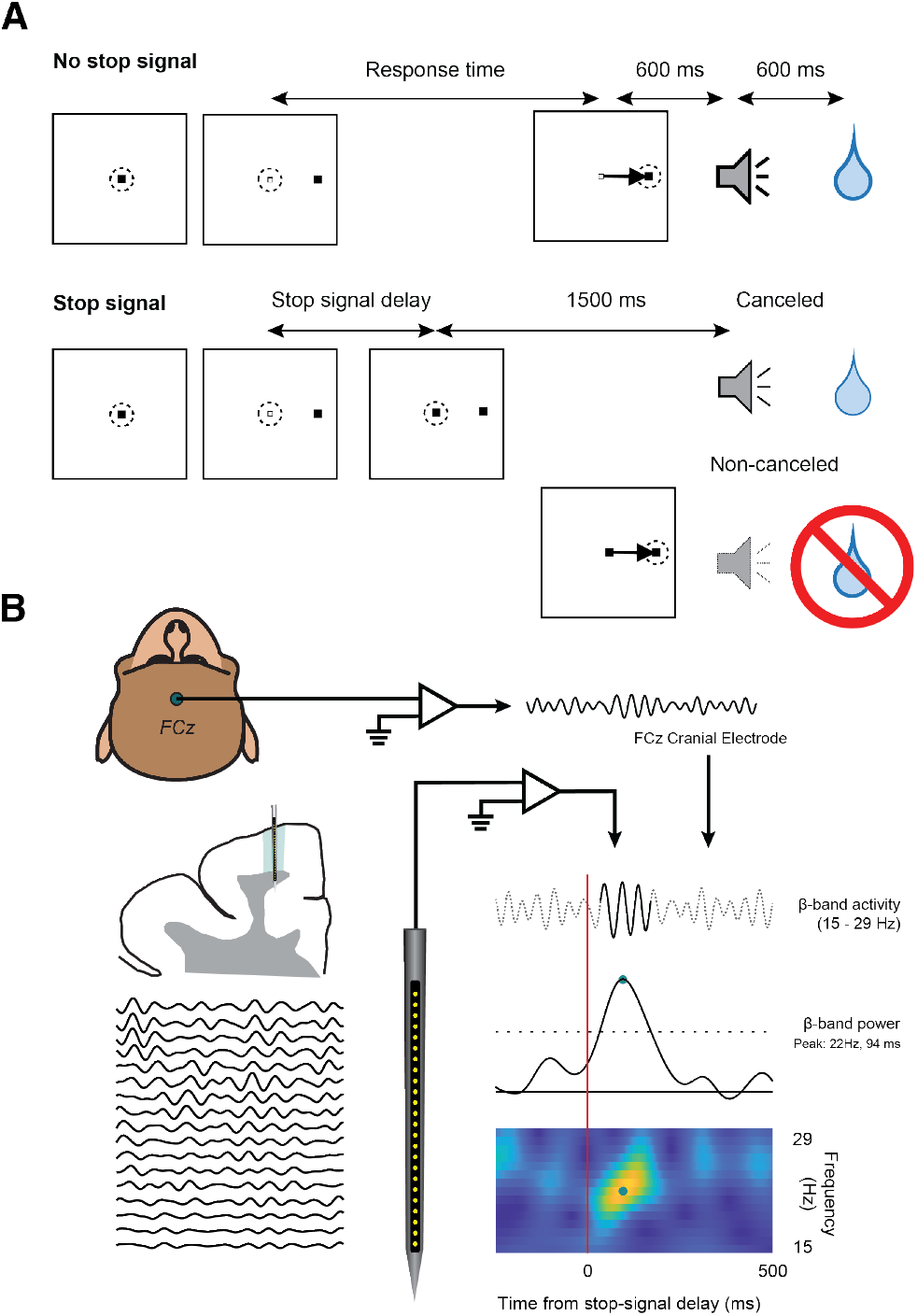
Experimental procedures. **A.** Saccade-countermanding task. Monkeys initiated trials by fixating on a central point. After a variable time, the center of the fixation point was extinguished. A peripheral target was presented simultaneously at one of two possible locations. On no-stop-signal trials, monkeys were required to shift their gaze to the target, whereupon after 600 ± 0 ms a high-pitched auditory feedback tone was delivered, and 600 ms later fluid reward was provided. On stop-signal trials (~40% of trials), after the target appeared the center of the fixation point was re-illuminated after a variable stop-signal delay, which instructed the monkey to cancel the saccade in which case the same high-pitched tone was presented after a 1,500 ± 0 ms hold time followed, after 600 ± 0 ms by fluid reward. The stop-signal delay was adjusted such that monkeys successfully canceled the saccade in ~50% of trials. In the remaining trials, monkeys made non-canceled errors which were followed after 600 ± 0 ms by a low-pitched tone, and no reward was delivered. Monkeys could not initiate trials earlier after errors. **B**. Neurophysiology methods. Electroencephalogram (EEG) was recorded with leads placed on the cranial surface over the medial frontal cortex at the location analogous to FCz in humans. Local field potentials were recorded from all channels from an electrode placed in the Supplementary Eye Field. For each session, raw data was bandpass filtered between 15 and 29 Hz and sampled from −1000 to +2500 ms relative to target presentation, saccade initiation, and stop-signal presentation. To detect β-bursts the signal in each interval for each trial was convolved with a complex Morlet wavelet. Time-frequency power estimates were extracted by calculating the squared magnitude of the complex wavelet-convolved data. Individual β-bursts were defined as local maxima in the trial-by-trial time-frequency power matrix, for which the power exceeded a threshold of 6-times the median power of the entire time-frequency power matrix for the electrode. An example burst is shown in the time-frequency plot.

Cortical local field potentials were sampled across 509 contacts (Eu: 217 contacts, X: 292 contacts), across 29 sessions (Eu: 12 sessions; X: 17 sessions), from electrodes placed in Supplementary Eye Field. In 16 of these sessions (Eu: 6; X: 10), penetrations were made perpendicular to the cortical surface. This allowed for the assignment of activity to specific cortical layers. Simultaneous to these recordings within the cortex, electroencephalogram (EEG) voltage was recorded with a lead placed on the cranial surface over the medial frontal cortex (**Fig. 1B**). The placement of these electrodes was analogous to FCz in humans. Like previous observations in EEG, the overall prevalence of cortical β-bursts was low during both baseline (−400 to −200 ms pre-target, 21.9 ± 0.6% across all sessions) and task-relevant (0 to 200 ms post-target, 22.8 ± 0.6% across all sessions) periods. Our first original finding is that the proportion of β-bursts observed within the cortex (21.9 ± 0.6%) mirrored those observed in EEG recorded on the cranial surface (baseline: 21.5 ± 0.9%, target: 27.9 ± 1.4%) during the baseline period of valid trials.

### Functional properties of intracortical β-bursts during executive control

Previous work has postulated that an increased incidence of β-bursts during the stopping period on canceled trials suggests a possible contribution to action-stopping. However, their role in error monitoring, overall sparsity, and prevalence in trials in which movements were executed warrants wider consideration of alternative functional contributions. Here, we more widely consider the pattern of medial frontal β-bursts across several task epochs reflecting different aspects of cognitive control (**Fig. 2A, Supplementary Fig. 2A**). We observe that the incidence of β-bursts clearly fluctuates over different task epochs and trial types, demonstrating notable variation in their functional contributions (Two-way repeated measures ANOVA, trial type x epoch: F (3.212, 1631.471) = 243.221, p < 0.001).

**Fig. 2.**
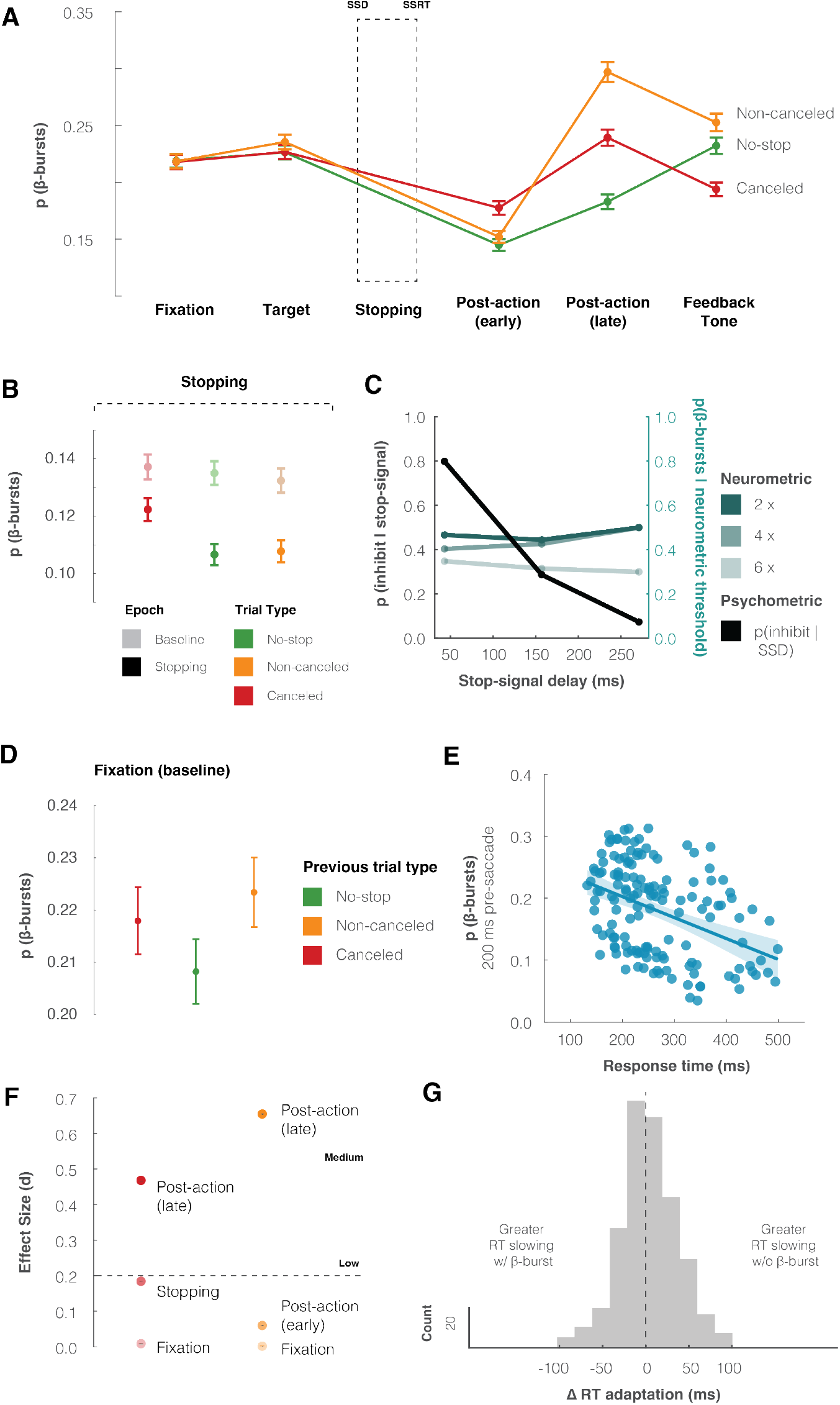
Functional properties of intracranial β-bursts. **A**. Mean ± SEM of the probability of LFP β-bursts across different epochs on canceled (red), non-canceled (yellow), and no-stop (green) trials. LFP β-bursts were more common in stop-signal trials and most common after errors. **B.** Mean ± SEM of LFP β-bursts observed during the STOP process interval between actual or scheduled SSD and SSRT during no-stop, non-canceled, canceled trials compared with an equivalent interval during foreperiod before target presentation (unsaturated). LFP β-bursts are more common in the foreperiod but observed in <15% of trials with slightly but significantly higher incidence when saccades are inhibited. **C**. Mismatch between p(inhibition) across SSD (black, left ordinate scale) for a representative session and neurometric functions across SSD (right ordinate scale) of p(β-burst) with median detection thresholds of 2x (darkest), 4x (intermediate), or 6x (lightest) baseline value. In no sessions were LFP β-bursts observed commonly enough to account for response inhibition. **D**. Mean ± SEM of p(β-bursts) during the foreperiod in no-stop trials following canceled, non-canceled and no-stop trials. LFP β-bursts were more common after error trials. **E**. Relationship between mean p(β-burst) in 200 ms preceding saccade initiation and means of session-wise response time quantiles, collapsed across all sessions. Regression line with CI demonstrates significant negative association. **F**. Mean ± SEM of bootstrapped effect sizes comparing the proportion of bursts in no-stop trials to canceled and non-canceled trials during fixation, stopping, early post-action, and late post-action epochs. LFP β-bursts were most common after responses, and evidence was weak for a higher incidence of LFP β-bursts during response inhibition. **G**. Histogram of change in RT between successive non-canceled and no-stop trials when an LFP β-burst did or did not occur during the error period. Slowing or speeding of RT after errors was unrelated to the occurrence of LFP β-burst after errors.

We first aimed to resolve the extent to which MFC β-bursts contributed to the active and direct inhibition of a planned response. Intracortical β-bursts were significantly less common during the STOP process compared to a pre-target baseline (**Fig. 2B**; two-way repeated measures ANOVA, main-effect of epoch: F (1, 508) = 226.52, p < 0.001). In conjunction with this, although observed in equal incidence during the baseline, β-bursts were significantly reduced during the potential stopping interval on trials in which a movement was generated, but not in trials in which a movement was inhibited (two-way repeated measures ANOVA, interaction between epoch and trial type: F (1.794, 911.151) = 31.812, p < 0.001). Mirroring previous observations in EEG (Enz et al., 2021; Hannah et al., 2020; Jana et al., 2020; Wessel, 2020), we found β-bursts were significantly more common during the STOP period when a planned saccade was inhibited (canceled trials), compared to when an erroneous movement was generated (non-canceled, adjusted p < 0.001), and to a comparable period in correct trials (no-stop, adjusted p < 0.001) trials. Whilst β-bursts were more common when stopping was successful, the relationship between the occurrence of β-bursts and behavioral measures of stopping was weak and unreliable across monkeys (**Supplementary Table 1)**.

Although the proportion of β-bursts was significantly different between trial types, there was no reliable relationship across monkeys between features of the β-burst (onset, offset, duration, frequency, volume) and trial types (**Supplementary Table 2, Supplementary Fig. 2B**). In addition, the prevalence of β-bursts was unchanging across stop-signal delays and did not reflect changes in the probability of inhibiting a saccade. The response inhibition function shows the fraction of non-canceled trials increases as a function of stop signal delay, as movements become less likely to be canceled as movement preparation progresses (**Fig. 2C, black line**). In conjunction with the psychometric function, a neurometric function can also be derived. Previous work has achieved this by inspecting the single neuron discharges of movement-related neurons in motor structures, such as the frontal eye fields (FEF), and determining the probability of single neurons modulating within the stopping period. Just like the psychometric function, this neurometric function also increases as a function of stop-signal delay (Brown et al., 2008). To examine if this relationship occurs with β-bursts, we measured the number of β-bursts observed in the last 50 ms of SSRT at several thresholds for each channel, on canceled trials at each stop-signal delay. We then calculated the sum of their squared differences relative to the probability of inhibiting a response at each stop-signal delay (**Fig. 2C**). To generate a valid null distribution, we repeated this process, but this time shuffling the p(β-bursts) observed at each SSD, before calculating the sum of squared differences. Across all channels, the summed squared differences were not significantly different from the shuffled condition at the 6x median threshold (two-sample t-test: t (1016) = 0.101, p = 0.920). This conclusion did not depend on the β-burst measurement threshold (**Supplementary Fig. 2C**, two-sample t-test at each threshold, p > 0.05 for all thresholds after corrections for multiple comparisons). Thus, β-bursting during the stopping period is unrelated to stopping behavior.

Second, we then considered that bursts may instead reflect another function supporting the ability to successfully stop, rather than the process of response-inhibition itself. We approached this from two perspectives: (1) activity before stopping that may influence or bias the stop process, and (2) activity following response inhibition that maintains inhibition until the response is validated and the goal is achieved (until the feedback tone is elicited).

To address the first perspective, we looked at how baseline activity could proactively influence the stop process. To do this, we compared the proportion of β-bursts that occurred during the fixation period between those trials which would ultimately progress to become inhibited, those which would lead to the generation of a correct saccade, and those which resulted in the generation of an erroneous saccade. β-bursts were not distinguishable in the fixation period of canceled, non-canceled, and no-stop trials (**Fig. 2A,** fixation, Holm post-hoc test, adjusted p > 0.999 for all comparisons). This observation was reliable across monkeys (**Supplementary Fig. 2A**). Second, we then looked at how baseline p(bursts) varied with the outcome of the previous trial. We compared p(β-bursts) occurring in the pre-target period (−400 to −200 ms) in no-stop trials following no-stop, canceled, and non-canceled trials. We found a small, but significant, reduction in p(β-bursts) following no-stop trials, compared to non-canceled and canceled trials (**Fig 2D**, one-way repeated measure ANOVA, F (1.836, 932.657) = 14.043, p < 0.001). Finally, we considered the proposition that β-bursts may influence the go process (Muralidharan et al., 2022) by looking at the relationship between β-bursts in saccade trials (both no-stop and non-canceled) and their relationship with RT. To do this, we looked at the proportion of β-bursts that occurred in two periods: (1) baseline/fixation and (2) pre-saccade. Although we found no relationship between baseline p(β-bursts) and RT (R^2^ = 0.012, p = 0.161), we did observe that the p(β-bursts) within the 200 ms pre-saccade window were increasingly less common as RT’s became longer (**Fig. 2E;** R^2^ = 0.169, p < 10^−5^). These findings were consistent across monkeys (**Supplementary Fig. 2D**).

Concerning the second perspective, we then examined patterns of β-burst activity following the period of reactive inhibition (between SSD and SSRT). To successfully complete the trial and receive a juice reward, monkeys had to maintain fixation on the central stop-signal following successful inhibition until a tone was sounded. During this period on canceled trials, we observed an increase in β-burst incidence following successful inhibition, compared to when a movement was correctly generated (**Fig. 2A,** early post-action, Holm post-hoc test, adjusted p < 0.001). This elevated bursting rate was maintained until the period immediately preceding the feedback tone (**Fig. 2A**, late post-action, Holm post-hoc test, adjusted p < 0.001). To determine whether this activity was stopping specific, we compared p(β-bursts) in this period to that in the fixation period. Here, the behavior is the same as that following successful stopping: the monkey was maintaining fixation at the same central point and waiting for the presentation of the next task cue – the presentation of a target for the fixation period, and the sounding of a tone for the stopping period. Interestingly, we observed a differential pattern of activity. Whilst β-bursts were observed in equal proportions on canceled and no-stop trials during the fixation period (**Fig. 2A**; fixation, adjusted p > 0.999), they were significantly more common on canceled trials in the early and late post-action periods (**Fig. 2A,** early post-action, adjusted p < 0.001; late post-action).

We quantified the magnitude of the difference in the p(bursts) between canceled and no-stop trials by calculating the effect size at three key epochs: during fixation, during stopping, and during the late post-action period. Although there were no differences between trial types during fixation, this developed into a low effect size during the stopping period, and a medium effect size during the late post-action period (**Fig. 2F**, two-way repeated-measures ANOVA, epoch x control signal, F(1.924, 190.468) = 7349.496, p < 0.001). Interestingly, β-bursts were less common on canceled trials and more common on no-stop trials following the feedback tone (**Fig. 2A**, Holm post-hoc test, adjusted p < 0.001). The increase of β-bursts in the early and late post-action period was consistent across monkeys (**Supplementary Fig. 2A**).

In addition to response inhibition, the stop-signal task is useful for exploring performance monitoring as, by design, errors occur in ~50% of stop-signal trials. Using this task, we previously reported that EEG β-bursts increase following errors (Errington et al., 2020). These findings mirrored previous work demonstrating increased spiking activity in SEF, occurring after errors (Purcell et al., 2012; Sajad et al., 2019; Stuphorn et al., 2010). Interestingly, contrasting our previous findings from EEG, we did not observe a reliable difference in the incidence of β-bursts in the 300 ms period following the execution of an erroneous or correct saccade. Instead, we observed a clear increase in β-bursts on error trials in the late post-action period leading to the auditory feedback tone (**Fig. 2A**). Congruent with this, we found no reliable difference in the proportion of β-bursts between error and correct saccade trials in the early post-action period between monkeys (**Supplementary Fig. 2A**, early post-action period, adjusted p = 0.116), but found β-bursts were significantly more common for both following errors in late post-action period (**Supplementary Fig. 2A**, late post-action period, adjusted p < 0.001).

Much like we did for trials in which inhibition was successful, we quantified the magnitude of the difference in the p(bursts) between non-canceled and no-stop trials by calculating the effect size at three key epochs: during the fixation, early post-action, and late post-action periods. There were no notable differences between error and correct trials during both fixation and early-action periods. This developed into a medium-to-large effect size during the late post-action period (**Fig. 2F**, two-way repeated-measures ANOVA, epoch x control signal, F (1.924, 190.468) = 7349.496, p < 0.001). Interestingly, in contrast to canceled trials, β-bursts were equally as common on non-canceled trials and no-stop trials following the feedback tone (**Fig. 2A**, Holm post-hoc test, adjusted p < 0.001).

To examine if these signals may be used to recruit control in future trials, we determined whether the presence of a β-burst in the late post-action period on error trials (trial *n*) influenced the RT on a following no-stop trial (trial *n+1*). For each channel, we compared the change in RT from non-canceled error trials to no-stop trials, between error trials with a β-burst present and those without a β-burst present. Here, negative values represent post-error slowing and positive represents an increase in RT’s following errors. We then calculated the difference in the RT adaptation between those trials with a burst against those without a burst. This provides a measure of the difference in RT adaptation between burst/no-burst trials. Here positive values represent greater post-error slowing when a burst was observed. In our sample, no channels demonstrated a relationship between bursting and RT adaptation (**Fig. 2G**). Thus, β-bursting during the error period is unrelated to RT adaptation.

### Laminar patterns of β-band activity

To investigate the cortical mechanisms contributing to β-bursts, we examined their profile across cortical layers of SEF from 16 sessions for which we had confidence in the layer assignments (Monkey Eu, 6 sessions; Monkey X, 10 sessions). The details of this procedure have been previously described (Godlove et al., 2014; Ninomiya et al., 2015).

We first used power spectral density (PSD) to look at the relative power of frequencies across all trials, and across all cortical depths. Qualitatively, we replicate previous findings that β-activity was most powerful in the middle-to-lower cortical layers, around the L3/5 divide in SEF (**Supplementary Fig. 1B**). This was observed across both monkeys (**Supplementary Fig. 1B**). To look at functional differences, we then compared β-power across all cortical layers on canceled trials relative to latency-matched no-stop trials during the STOP process. We found that normalized β-power was significantly greater in L5 & L6 on canceled trials, but not in L2 & L3 (**Fig. 3A,** two-way mixed-measures ANOVA, trial type x cortical layer interaction: F (1, 247) = 6.633, p = 0.011). However, we note this effect was driven by one monkey (**Supplementary Fig. 3A**).

**Fig. 3.**
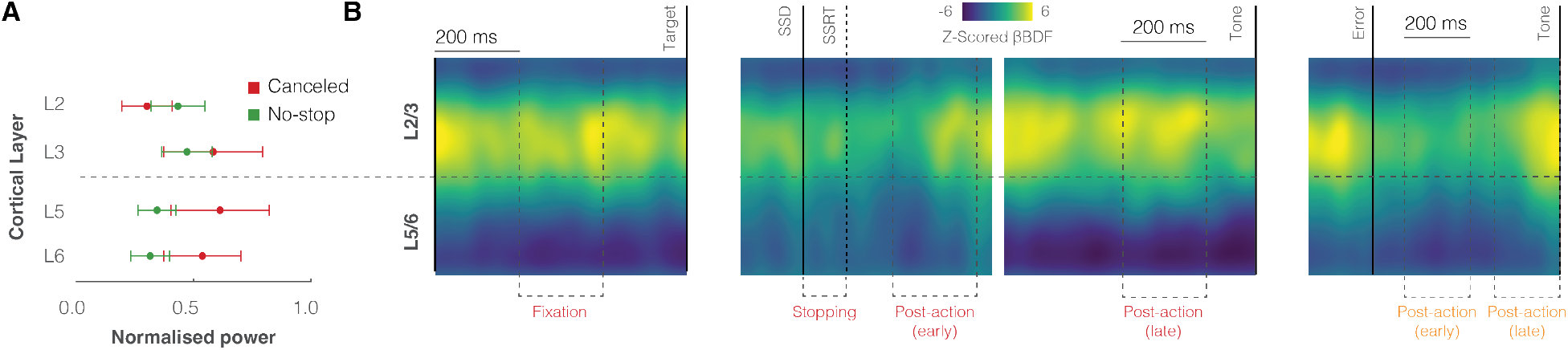
Laminar properties of intracranial β-bursts. **A**. Normalized power in each cortical layer during response inhibition for canceled and latency-matched no-stop trials. The elevation of LFP β power during response inhibition happened in the deep layers. Conventions as in Fig. 1. **B**. Heat maps of LFP β-burst incidence across time and cortical layers on canceled trials during foreperiod (left), stopping and early post-action (left-center), late post-action period (right-center), and error (right) epochs. Yellow represents a greater proportion of (β-bursts) at a given time and depth, relative to other times and depths.

We then progressed to look at the pattern of β-bursts, rather than power, through cortical layers across different functional epochs within our task. We compared the proportion of β-bursts observed in each layer during fixation, stopping, early, and late post-action periods on canceled trials, and the late post-action period on noncanceled trials. Although variable at the individual layer level between monkeys, we observed bursting was reliably more prevalent in L3 for both monkeys (**Supplementary Fig. 3B**, two-way mixed-measures ANOVA, main effect of cortical layer: F (3, 245) = 27.734, p < 0.001).

To examine the temporal profile of this activity, we generated time-depth β-burst density functions which allow us to see the pattern of β-bursts across cortical layers relative to different task events. Given the higher incidence of beta-bursts during these epochs, we examined the laminar pattern of bursts during fixation, stopping, and in the late post-action period on canceled trials, and the early and late post-action period on noncanceled error trials. For each session, this resulted in a β-burst density function over time and depth which was then z-scored relative to the mean value across all depths and during the entire epoch; higher Z-scores represent a greater proportion of β-bursts observed relative to other times and depths. Using this approach, we observed that β-bursts were qualitatively more prevalent in upper layers compared to lower layers during fixation, stopping, and errors (**Fig. 3B**). This effect was observed in both monkeys (**Supplementary Fig. 3C**).

### Weak association between intracortical and cranial β-bursts

After establishing the patterns of β-bursts activity within SEF, we then determined the extent to which this activity could contribute to β-bursts recorded on the cranial surface through several approaches. We first compared the pattern of bursts observed in the cortex with the pattern of bursts observed in the simultaneously recorded EEG signal (**Fig. 4A**). We quantified the proportion of bursts in each epoch in each electrode channel, across all channels in the upper and lower layers, and across all channels in the cortex. In the limit, EEG β-bursts could be generated by β-bursts observed simultaneously in all sampling channels across all cortical layers. Alternatively, EEG β-bursts could be generated by β-bursts observed simultaneously in some fraction of sampling channels across cortical layers. In the other limit, EEG β-bursts could appear sporadically with no relationship to β-bursts arising randomly in sampling channels across cortical layers.

**Fig. 4.**
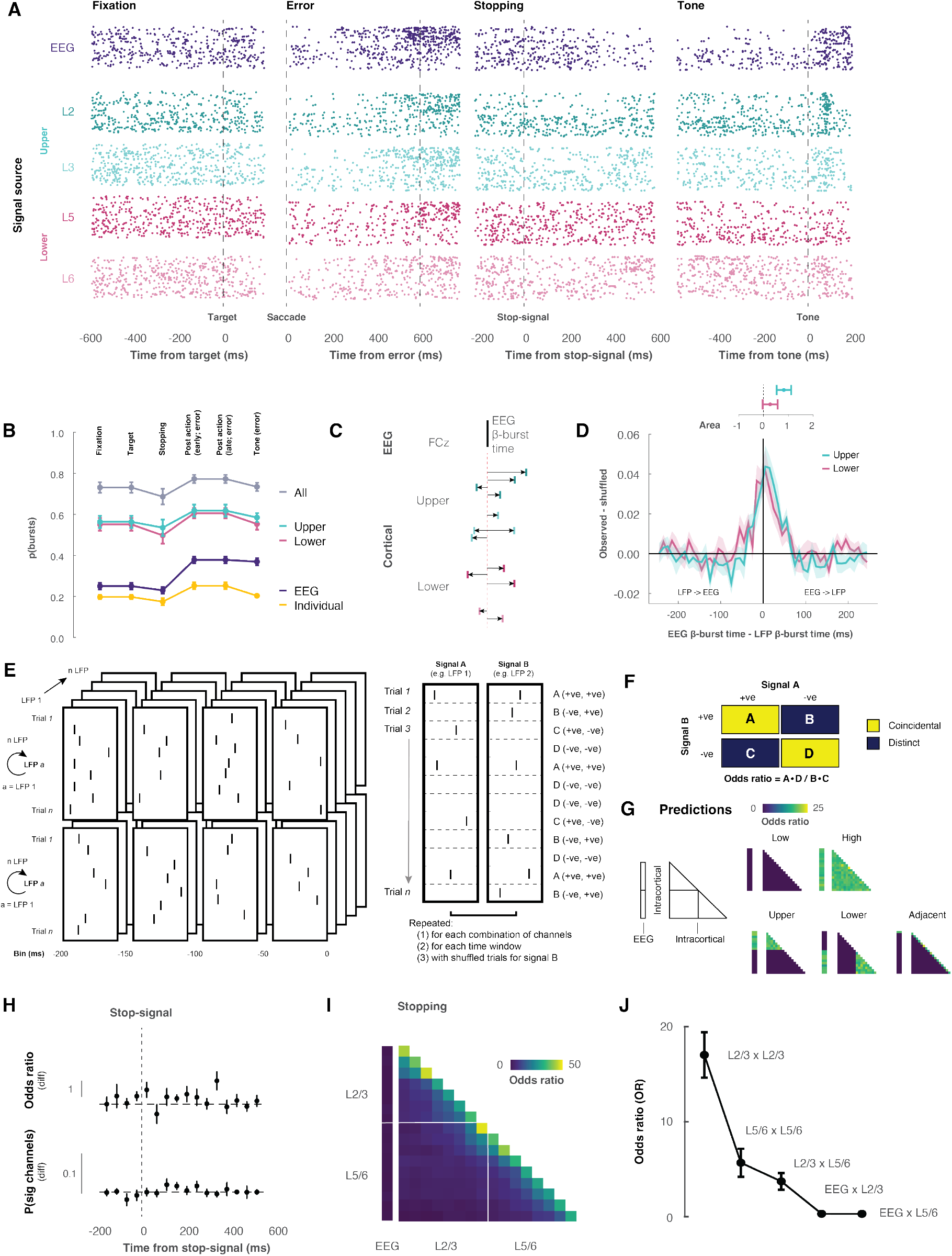
Comparing EEG and LFP β-bursts. **A.** With each row representing a trial from a representative session, each point in the raster marks the time EEG (purple) the LFP β-bursts in L2 (dark teal), L3 (light teal), L5 (dark magenta), L6 (light magenta) during fixation (left), error (left-center), stopping (right-center), and canceled late post-action (right) epochs. Note the lack of columnar synchrony with density after errors and with feedback. **B**. Mean ± SEM of p(β-burst) counted within epochs in individual LFP (gold) and EEG (purple) contacts and summed across contacts in L2/3 (teal), in L5/6 (magenta), and from L2 to L6 (grey). The number of LFP β-bursts within a cortical column exceeds the number of EEG β-bursts. **C**. Diagram of how the coincidence of EEG and LFP β-bursts was measured. **D**. Distribution after shuffle subtraction of shortest times between EEG β-bursts and LFP β-bursts detected in L2/3 (teal) and L5/6 (magenta) plotted with positive (negative) values for LFP β-bursts following (preceding) EEG β-bursts. Only ~5% of EEG β-bursts were associated more than chance with LFP β-bursts, and this association was observed only within ±50 ms of the EEG β-burst. Mean ± SEM area under the distributions across sessions, plotted above, indicates a weak tendency for LFP β-bursts in L2/3 but not L5/6 to follow EEG β-bursts. **E.** Diagram of how odds ratio (OR) of presence or absence of EEG and LFP β-bursts in non-overlapping 50 ms intervals surrounding task events. The presence or absence of β-bursts was counted pairwise across every EEG and LFP channel for each interval of all trials. **F**. Odds ratios were calculated from the counts of the four possible combinations of presence or absence of β-bursts in each pair of channels were tabulated: β-burst present in both (A), in one channel but not the other (B, C), and β-burst absent in both (D). **G.** Predicted odds ratios were calculated in the matrix of EEG and LFP sampling contacts for simulations of coincident presence or absence of β-bursts occurring rarely (low) or frequently (high) across all channels or occurring frequently only in the upper, lower, or adjacent channels. **H**. Mean ± SEM of the shuffle-corrected odds ratio of detecting an LFP β-burst ±50 ms from the occurrence of an EEG β-burst (above) and of the probability of detected significant odds ratio > 1 (below) during response inhibition. Coincident presence or absence of β-bursts across EEG and LFP channels was vanishingly rare. **I**. Color map of odds ratios of LFP and EEG β-bursts shows association in adjacent but not distant contacts. **J**. Mean ± SEM odds ratios of coincidences of β-bursts occurring within L2/3 LFP channels, within L5/6 LFP channels, across L2/3 and L5/6 LFP, and across EEG and LFP in L2/3 and L5/6. The odds ratio scaled with prevalence and was effectively nil across EEG and LFP channels.

By simply counting the number of β-bursts in the EEG electrode and each LFP channel in each task epoch, we found that on average they were infrequent on any individual LFP channel and only slightly more frequent in the EEG after saccades (**Fig. 4B; Supplementary Fig. 4A**). When summed within L2/3 and L5/6 or across all layers, the incidence of β-bursts was naturally higher. The sum of β-bursts within L2/3 and L5/6 or across all LFP channels exceeds by 3 or 4 times, the number of β-bursts in the EEG channel. This reveals unexpected complexity in the biophysical relationship between LFP and EEG β-bursts.

We then quantified the temporal relationship between LFP and EEG β-bursts (**Fig. 4C**) and found that β-bursts were significantly more likely in both L2/3 and L5/6 in the ±50 ms interval around the EEG β-burst (**Fig. 4D, main**). LFP β-bursts were more likely to occur after EEG bursts, than before (**Fig. 4D, inset;** one-sample t-test (two-tailed), t(15) = 2.167, p = 0.047). Although we observed between-monkey differences (**Supplementary Fig. 4B**), we reliably observe this finding for bursts in upper relative to lower cortical layers (Lower: one-sample t-test (two-tailed), t(15) = 0.904, p = 0.380; Upper: one-sample-test (two-tailed), t(15) = 2.821, p = 0.012).

Finally, having determined the ±50 ms window of association of EEG and LFP β-bursts, we quantified the coincident presence or absence of EEG and LFP β-bursts. This was achieved by identifying bursts trial-by-trial in non-overlapping 50 ms windows around the event of interest for every LFP contact within the cortex and the EEG signal (**Fig. 4E. left**). For each possible pair of contacts trial-by-trial we counted how often a β-burst co-occurred, occurred in one channel or the other, or neither channel (**Fig. 4E, right**) and calculated the odds ratio (**Fig. 4F**). This odds ratio provides a measure of association between observing a burst in one channel and another burst in a separate channel. To test alternative hypotheses about β-burst production, we simulated low and high coincidence between all contacts, only between contacts in the upper or lower layers, and only between adjacent contacts (**Fig. 4G**).

Relative to shuffled control, no periods of significant association between the proportion of EEG and LFP β-bursts were observed in any task epoch (**Fig. 4H; Supplementary Fig. 4C, D**). We also considered the intra-contact similarity to probe the coincidence of bursts between and within cortical layers (**Fig. 4I; Supplementary Fig. 4E**). β-bursts coincided significantly more on adjacent contacts within layers than across layers (**Fig. 4J; Supplementary Fig. 4F**, two-way independent-groups ANOVA, main effect of laminar comparison: F (3, 300) = 103.151, p < 0.001). This relationship was more pronounced in upper cortical layers compared to lower cortical layers (adjusted p < 0.001) and did not vary between epochs (interaction between epoch and laminar comparison: F (3, 300) = 0.102, p > 0.999). Finally, the OR of the association between LFP and EEG β-bursts was significantly smaller than the OR of the association between β-bursts within the cortex (**Fig. 4J**, adjusted p < 0.001 for all comparisons). These results demonstrate that β-bursts are not produced concomitantly within a cortical column but instead arise at low and stochastic rates relatively independently across cortical depth.

## DISCUSSION

### Functional properties of intracranial medial frontal β-bursts

Previous work in humans (Wessel, 2020) and macaques (Errington et al., 2020) has described a higher prevalence of EEG β-bursts recorded over the medial frontal cortex (MFC) during response inhibition. Like other reports of EEG β-bursts in the frontal cortex (REF), the incidence of these bursts was low and their association with stopping behaviors was weak. This lack of causal efficacy is important as it limits the potential theoretical and practical applicability of such signals.

We proposed several reasons why EEG β-bursts may only weakly index stopping. First, the rarity of β-bursts may result from the low signal-to-noise ratio (SNR) of EEG recordings. We addressed this directly by recording directly from the cortex, adjacent to where these signals are recorded from, using microelectrode arrays. As we record directly from the cortex, we reduce any filtering of neural activity by the skull, muscles, and other tissue, effectively allowing us to increase the SNR. In our intracortical sample, we observed that β-bursts became less common as a saccade was being prepared, consistent with interpretations proposing β-activity acts as a brake on movement or decisions (Muralidharan et al., 2022). Although we observed that β-bursts were less common prior to a longer-latency saccade being generated, we note that this may be resultant of contamination of a marked visual transient β-burst response at earlier response latencies. Importantly, although β-bursts were more prevalent during the period in which a saccade was inhibited, they were infrequent, and their properties did not account for the likelihood of canceling a planned response. As such, despite improving the SNR, we did not see any increase in the incidence of β-bursts or any improvement in their relation to stopping behaviors.

This finding demonstrates that β-bursts within the Supplementary Eye Field cannot act as a causal mechanism for reactive response inhibition. Instead, we consider that β-bursts may best reflect proactive executive control processes. First, we observed an increased likelihood of β-bursts following errors. Although these bursts were not associated with post-error adjustments in behavior, these findings mirror previous observations of error-related activity in the medial frontal cortex (Emeric et al., 2010; Purcell et al., 2012; Sajad et al., 2019; Stuphorn et al., 2000). Interestingly, β-bursts were more frequent well after the error, just before the feedback tone. Second, we observed a prolonged, higher incidence of β-bursts after successful stopping. After successfully inhibiting a plan to generate a movement, our task required monkeys to maintain fixation until a feedback tone sounded. Hence, β-bursts during this post-stopping period could be interpreted in the context of previous findings implicating β-activity in “clearing out” working memory (Lundqvist et al., 2016) and blocking distractions (Castiglione et al., 2019; Hanslmayr et al., 2014). These processes may allow the maintenance of task goals (e.g., sustain unblinking fixation) that will result in reward (Sajad et al., In Press). Collectively, these observations both provide evidence that SEF signals are most prominent following an action, supporting previous theories of medial frontal cortex function (Bonini et al., 2014; Passingham et al., 2010; Rushworth et al., 2004; Schall et al., 2002).

Interestingly, these observations also highlight a clear distinction between human and macaque models of stopping, with particular regard to the role of the medial frontal cortex. Whilst human studies emphasize that the pre-SMA is active during response inhibition and can contribute to the interruption of response preparation (Diesburg and Wessel, 2021; Rae et al., 2014; Sharp et al., 2010), neurophysiological evidence obtained in the medial frontal cortex of macaques demonstrate that neurons in this area modulate too late to directly contribute to stopping (Scangos and Stuphorn, 2010; Stuphorn et al., 2010). In our introduction, we proposed that this divergent evidence between macaques and humans may be resultant of differences intrinsic to the methodologies and analyses used.

We attempted to address this by mirroring recording and analysis techniques used in humans and combining them with intracortical recordings afforded by macaque neurophysiology. Through our previous work, we demonstrated that β-bursts in EEG had functionally similar properties to those in humans; their incidence decreased as a movement was being prepared and was slightly more common during response inhibition (Errington et al., 2020). Interestingly and unexpectedly, however, although we mirror the functional properties of EEG β-bursts in our intracortical signals, there was no association between the presence of a burst within the cortex and one within EEG. This is an important null result, the confidence in which is engendered by other clear, specific, and positive associations observed between neural signals in SEF with overlying EEG (Sajad et al. 2019, 2022; Herrera 2020) and other examples of dissociated SEF LFP vs. EEG during performance monitoring (Westerberg et al., 2020). However, further investigation is needed to determine the extent to which EEG β-burst production sampled over other cortical areas is related to LFP β-bursts in the underlying dipole sources. We also, however, note one important confound: human studies classically employ a manual countermanding task, in which participants move a joystick or press a button, whilst macaque studies (including ours) employ a saccade countermanding task. Although pre-SMA and SEF have similar functions for their respective effectors (Fujii et al., 2002), and models of response inhibition are applicable across different modalities (Boucher et al., 2007b; Logan and Irwin, 2000), there may be significant anatomical and functional differences in the stopping circuitry between these two which lead to variations in the contribution of different brain areas to the EEG signal. This warrants further investigation.

### Origin of β-burst in agranular neocortex and EEG

Bioelectric potentials have practical and clinical applications when their generators are known. For example, the electrocardiogram is useful in medicine because the physiological process associated with each phase of polarization is understood. Likewise, the electroretinogram is useful because the cell layers associated with each polarization are understood. In contrast, signals recorded over the scalp indexing cognitive operations will have limited utility until their neural generators are known. This work adds to a growing literature addressing this issue by simultaneously recording signals in the brain and on the cranial surface (Bimbi et al., 2018; Cohen et al., 2009; Cosman et al., 2018; Heitz et al., 2010; Herrera et al., 2022; Purcell et al., 2013; Sandhaeger et al., 2019; Shin et al., 2017; Westerberg et al., 2020; Westerberg et al., 2022; Whittingstall and Logothetis, 2009).

By combining the laminar profile of β-bursts described in this study with knowledge of the anatomical and histological properties of SEF, we can gain further insight into how β-bursts can arise and how they relate to β-bursts recorded on the cranial surface. A recent model using 100 multi-compartment pyramidal cells in layer 3, another 100 in layer 5 with 35 inhibitory neurons in layer 3 and layer 5 showed how β-bursts can be produced via a transient dipole reversal caused by the interaction between a prolonged (>100 ms) excitatory synaptic drive in the basal dendrites of pyramidal neurons that interacts with a transient pulse of excitatory drive in the apical dendrites persisting for at least 50 ms (Sherman et al., 2016; Shin et al., 2017). Consistent with this, we found β-bursts were more prevalent in upper cortical layers across all task epochs. The core mechanism embodied by the model of synchronous patterns of basal and apical dendritic events has been proposed in other contexts such as error detection (Cohen, 2014), and another recent biophysical model of layer 5 pyramidal neurons highlighted the contributions of apical dendrite calcium spikes (Herrera et al., 2020). The interpretation of the origin and role of β-bursts is enhanced by converging observations about the properties of neurons across the layers of SEF (Sajad et al., 2019)(Sajad et al. 2022), which demonstrate contributions to error detection, reinforcement expectation, response conflict, event timing, and working memory.

Interestingly, we found that LFP β-burst presence was not synchronized across the cortical column but instead coincided only on adjacent LFP channels. The spacing of the linear electrode contacts (150 μm) indicates that, assuming an isotropic medium, intracranial β-bursts occupy a volume no more than ~0.014 mm^3^. These observations offer useful constraints on biophysical models of β-burst production (Sherman et al. 2016).

### Cautionary considerations of β-burst activity

Sampling field potentials from within the cerebral cortex was supposed to improve the poor signal-to-noise ratio (SNR) of noninvasive EEG recordings. Nevertheless, we observed β-bursts as rarely in intracortical LFP as in cranial EEG recordings. Combining simultaneously recorded EEG and LFP signals allowed a direct comparison of β-bursts in two techniques thought to differ in SNR. Interestingly, at the individual channel level, we observed that β-bursts were as rare in intracranial LFPs as they were in EEG signals. Despite being similar in incidence, LFP β-bursts were not coincident with β-bursts observed in EEG. Furthermore, the incidence of β-bursts observed within MFC parallels that observed in human intracranial recordings (Diesburg et al., 2021; Mosher et al., 2021; Yu et al., 2021).

These results urge caution when interpreting β-bursts across cortical layers. By definition, β-bursts are detected when β-band power crosses a pre-defined threshold, based on the average overall power at a site. This approach may result in a lesser probability for a burst to be detected at sites with greater β-power. In this study we took two approaches to this potential problem, considering the threshold as a function of either the power at the given contact or the power across the entire site (all contacts within the cortex in the penetration). We believe taking the power across the entire site results in more balanced and fair opportunities for a burst to be captured and have presented data from that approach in this manuscript. Nevertheless, both approaches yielded qualitatively similar patterns that allow for the preposition of ideas on the functional architecture of β-burst signaling in the medial frontal cortex.

### Conclusion

These results demonstrate that β-bursts are about as rare in the cerebral cortex as they are on the cranial surface and exhibit no more reliable association with response inhibition than that previously reported for EEG β-bursts. The findings demonstrate that β-bursts are incapable of mediating any mechanism of response inhibition. Nevertheless, β-bursts in MFC may index other processes underlying executive control.

## MATERIALS & METHODS

### Experimental models and subject details

Data was collected from one male bonnet macaque (Eu, *Macaca Radiata*, 8.8 kg) and one female rhesus macaque (X, *Macaca Mulatta*, 6.0 kg) performing a countermanding task (Godlove et al., 2014; Hanes and Schall, 1995). All procedures were approved by the Vanderbilt Institutional Animal Care and Use Committee in accordance with the United States Department of Agriculture and Public Health Service Policy on Humane Care and Use of Laboratory Animals.

### Animal care and surgical procedures

Surgical details have been described previously (Godlove et al., 2011). Briefly, magnetic resonance images (MRIs) were acquired with a Philips Intera Achieva 3T scanner using SENSE Flex-S surface coils placed above or below the animal’s head. T1-weighted gradient-echo structural images were obtained with a 3D turbo field echo anatomical sequence (TR = 8.729 ms; 130 slices, 0.70 mm thickness). These images were used to ensure that Cilux recording chambers were placed in the correct area (Crist Instruments). Chambers were implanted normal to the cortex (Monkey Eu: 17°; Monkey X: 9°; relative to stereotaxic vertical) centered on the midline, 30 mm (Monkey Eu) and 28mm (Monkey X) anterior to the interaural line.

### Cortical mapping and electrode placement

Chambers implanted over the medial frontal cortex were mapped using tungsten microelectrodes (2-4 MΩ, FHC, Bowdoin, ME) to apply 200 ms trains of biphasic micro-stimulation (333 Hz, 200 *μ*s pulse width). The SEF was identified as the area from which saccades could be elicited using < 50 *μ*A of current (Martinez-Trujillo et al., 2004; Schall, 1991; Thompson et al., 1996). In both monkeys, the SEF chamber was placed over the left hemisphere.

A total of five penetrations were made into the cortex; two in monkey Eu and three in monkey X. Three of these penetrations were perpendicular to the cortex. In monkey Eu, the perpendicular penetrations sampled activity at site P1, located 5 mm lateral to the midline and 31 mm anterior to the interaural line. In monkey X, the perpendicular penetrations sampled activity at sites P2 and P3, located 5 mm lateral to the midline and 29 and 30 mm anterior to the interaural line, respectively. However, during the mapping of the bank of the cortical medial wall, we noted both monkeys had chambers placed ~1 mm to the right respective to the midline of the brain. This was confirmed through co-registered CT/MRI data. Subsequently, the stereotaxic estimate placed the electrodes at 4mm lateral to the cortical midline as opposed to the skull-based stereotaxic midline.

### Data acquisition

Spiking activity and local field potentials were recorded from five sites within the SEF using a 24-channel Plexon U-probe (Dallas, TX) with 150 *μ*m interelectrode spacing allowing sampling from all layers. Penetrations in three of these sites were perpendicular to the cortex. The U-probes were 100 mm in length with 30 mm reinforced tubing, 210 *μ*m probe diameter, 30° tip angle, and 500 *μ*m between the tip and first contact. Contacts were referenced to the probe shaft and grounded to the headpost. We used custom-built guide tubes consisting of 26-gauge polyether ether ketone (PEEK) tubing (Plastics One, Roanoke, VA) cut to length and glued into 19-gauge stainless steel hypodermic tubing (Small Parts Inc., Logansport, IN). This tubing had been cut to length, deburred, and polished so that they effectively support the U-probes as they penetrated the dura and entered the cortex. The stainless-steel guide tube provided mechanical support, while the PEEK tubing electrically insulated the shaft of the U-probe, and provided an inert, low-friction interface that aided in loading and penetration.

Microdrive adapters were fit to recording chambers with < 400 *μ*m of tolerance and locked in place at a single radial orientation (Crist Instruments, Hagerstown, MD). After setting up hydraulic microdrives (FHC, Bowdoin, ME) on these adapters, pivot points were locked in place using a custom mechanical clamp. Neither guide tubes nor U-probes were removed from the microdrives once recording commenced within a single monkey. These methods ensured that we were able to sample neural activity from precisely the same location relative to the chamber on repeated sessions.

Electrophysiology data were processed with unity-gain high-input impedance head stages (HST/32o25-36P-TR, Plexon). All data were streamed to a single data acquisition system (MAP, Plexon, Dallas, TX). Time stamps of trial events were recorded at 500 Hz. Eye position data were streamed to the Plexon computer at 1 kHz using an EyeLink 1000 infrared eye-tracking system (SR Research, Kanata, Ontario, Canada).

### Stop-signal task

The saccade stop-signal (countermanding) task utilized in this study has been widely used previously (Cabel et al., 2000; Colonius et al., 2001; Godlove and Schall, 2016; Hanes and Carpenter, 1999; Hanes and Schall, 1995; Kornylo et al., 2003; Morein-Zamir and Kingstone, 2006; Thakkar et al., 2011; Thakkar et al., 2015; Verbruggen et al., 2019; Walton and Gandhi, 2006; Wattiez et al., 2016). Briefly, trials were initiated when monkeys fixated on a central point. Following a variable period, the center of the fixation point was removed leaving an outline. At this point, a peripheral target was presented simultaneously on either the left or right hand of the screen. In this study, one target location was associated with a larger magnitude of fluid reward. The lower magnitude reward ranged from 0 to 50% of the higher magnitude reward amount. This proportion was adjusted to encourage the monkey to continue responding to both targets. The stimulus-response mapping of location-to-high reward changed across blocks of trials. Block length was adjusted to maintain performance at both targets, with the number of trials in each block determined by the number of correct trials performed. In most sessions, the block length was set at 10 to 30 correct trials. Erroneous responses led to repetitions of a target location, ensuring that monkeys did not neglect low-reward targets in favor of high-reward targets – a phenomenon demonstrated in previous implementations of asymmetrically rewarded tasks (Kawagoe et al., 1998).

In most of the trials, the monkey was required to make an eye movement toward this target (no-stop trials). However, in a proportion of trials the center of the fixation point was re-illuminated (stop-signal trials); this stop-signal appeared at a variable time after the target had appeared (stop-signal delay; SSDs). An initial set of SSDs, separated by either 40 or 60 ms, were selected for each recording session. The delay was then manipulated through an adaptive staircasing procedure in which stopping difficulty was based on performance. When a subject failed to inhibit a response, the SSD was decreased by a random step to increase the likelihood of success on the next stop trial. Similarly, when subjects were successful in their inhibition, the SSD was increased to reduce the likelihood of success on the next stop trial. This procedure was employed to ensure that subjects failed to inhibit action on approximately 50% of all stop-signal trials. On no-stop trials, the monkey was rewarded for making a saccade to the target. In stop-signal trials, the monkey was rewarded for withholding the saccade and maintaining fixation on the fixation spot. Following a correct response, an auditory tone was sounded 600ms later, and followed by a high or low fluid reward, depending on the stimulus-response mapping.

### Data collection protocol

An identical daily recording protocol across monkeys and sessions was carried out. In each session, the monkey sat in an enclosed primate chair with their head restrained 45cm from a CRT monitor (Dell P1130, background luminance of 0.10 cd/m2). The monitor had a refresh rate of 70Hz, and the screen subtended 46 deg x 36 deg of the visual angle. Eye position data were collected at 1 kHz using an EyeLink 1000 infrared eye-tracking system (SR Research, Kanata, Ontario, Canada). This was streamed to a single data acquisition system (MAP, Plexon, Dallas, TX) and amalgamated with other behavioral and neurophysiological data. After advancing the electrode array to the desired depth, they were left for 3 to 4 hours until recordings stabilized across contacts. This led to consistently stable recordings. Once these recordings stabilized, an hour of resting-state activity in near-total darkness was recorded. This was followed by the passive presentation of visual flashes followed by periods of total darkness in alternating blocks. Finally, the monkey then performed approximately 2000 to 3000 trials of the saccade countermanding (stop-signal) task.

### Quantification and Statistical Analysis

#### Cortical depth assignment

The retrospective depth of the electrode array relative to grey matter was assessed through the alignment of several physiological measures. Firstly, the pulse artifact was observed on a superficial channel which indicated where the electrode was in contact with either the dura mater or epidural saline in the recording chamber; these pulsated visibly in synchronization with the heartbeat. Secondly, a marked increase of power in the gamma frequency range (40-80 Hz) was observed at several electrode contacts, across all sessions. Previous literature has demonstrated elevated gamma power in superficial and middle layers relative to deeper layers (Bastos et al., 2018; Godlove et al., 2014; Maier et al., 2010; Ninomiya et al., 2015; Smith and Sommer, 2013; Westerberg et al., 2019; Xing et al., 2012). Thirdly, an automated depth alignment procedure was employed which maximized the similarity of CSD profiles evoked by passive visual stimulation between sessions (Godlove et al., 2014).

### LFP processing and β-burst detection

For each session, raw data was extracted from the electrode. This signal was then bandpass filtered between 15 and 29 Hz. This signal was then epoched from −1000 ms to 2500 ms relative to multiple key events in a trial, including target onset, saccade, and stop-signal presentation. β-burst detection was performed as previously described (Shin et al., 2017; Wessel, 2020). The description is adapted from therein. We then convolved the epoched signal for each trial with a complex Morlet wavelet of the form:

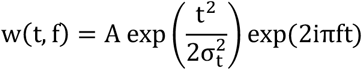

with 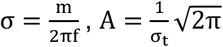, and m = 7 (cycles) for each of the 15 evenly spaced frequencies spanning the β-band (15 – 29 Hz). Time-frequency power estimates were extracted by calculating the squared magnitude of the complex wavelet-convolved data. Individual β-bursts were defined as local maxima in the trial-by-trial band time-frequency power matrix, for which the power exceeded a threshold of 6-times the median power of the entire time-frequency power matrix for the electrode. To compute the burst % across trials, we binary-coded the time of the peak β-amplitude. A β-burst density function was generated by convolving the binary-coded array of β-burst activity with a Gaussian function of the form:

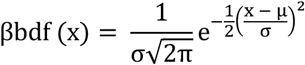

where μ = 00.0 ms, and σ = 22.5 ms. These values represent the time required for half a cycle at the median β-frequency.

### General analysis considerations

All statistical comparisons were conducted in JASP 0.16.2. When conducting ANOVAs, Mauchly’s test of sphericity was used to assess sphericity. Greenhouse-Geisser corrections were employed when this assumption was violated. When multiple comparisons were made, p-values were adjusted using the Holm method. When binning data, we simply recorded whether one or more bursts occurred within the bin; and did not code the number of bursts within a given bin. For all analyses, except neurometric, we defined bursts as periods in which the LFP power was greater than 6-times the median power of the entire electrode. This parameter was varied between 1 x and 10 x the median power in the neurometric approach.

### Bayesian modeling of stop-signal performance

As performance on the stop-signal task can be considered as the outcome of a race between a GO and STOP process, then a stop-signal reaction time (SSRT) can be calculated (Logan and Cowan, 1984). This value can be considered as the latency of the inhibitory process that interrupts movement preparation. Stop-signal reaction time was estimated using a Bayesian parametric approach (Matzke et al., 2013a; Matzke et al., 2013b). Compared to classical methods of calculating SSRT (integration-weighted method, Logan and Cowan (1984)), this approach allows for a distribution of SSRT to be derived by using the distribution of reaction times on no-stop trials, and by considering reaction times on non-canceled trials as a censored no-stop response time (RT) distribution. Furthermore, this model also allows for the estimation of the probability of trigger failures for a given session (Matzke et al., 2017). Individual parameters were estimated for each session. The priors were bounded uniform distributions (*μ*_*Go*_, *μ_Stop_: U* (0.001, 1000); *σ*_*Go*_, *σ_Stop_: U* (1, 500) *τ*_*Go,*_ *τ_Stop_: U* (1, 500); pTF: *U* (0,1)). The posterior distributions were estimated using Metropolis-within-Gibbs sampling and we ran multiple (3) chains. We ran the model for 5000 samples with a thinning of 5.

### Examining the functional properties of medial frontal β-bursts

We used a repeated-measures ANOVA to compare bursts during the stop period (from SSD to SSRT) and baseline period (−200-SSRT to −200 ms, pre-target) across all canceled, non-canceled, and no-stop trials for each contact within the cortex (n = 509). We observed the assumption of sphericity was violated (Mauchly’s test of sphericity, p <.05), so used Greenhouse-Geisser corrections when considering the outcome of our ANOVA analysis. P-values for our post-hoc comparisons were Holm adjusted for comparing a family of 15 (3 trial types, 2 periods).

We also considered how the properties of β-bursts may relate to behavioral measures of stopping. For this, we determined several properties of bursts that occurred in the STOP period: from SSD to SSRT. For no-stop and non-canceled trials, we assigned a pseudo-SSD based on the SSD of the previous trial. P(Burst) was calculated as the proportion of trials in which a burst was observed during the period of interest. Onset and offset were determined as the time at which the activity crossed the defined power threshold, relative to the time of the identified peak. The duration was determined as the time from onset to offset. Frequency was identified as the frequency at which the β-burst peak was located. We also determined the range of frequencies a burst occupied, by finding the frequencies closest to the burst peak at which the power crossed the defined threshold. Volume was calculated as the duration of the burst, multiplied by the amplitude, multiplied by the range of frequencies that it covered. As the behavioral measure was estimated on a session-by-session basis, we averaged the properties of all bursts observed within a session. To examine the relationship between behavioral variables of stopping (such as the estimated mean SSRT, estimated variance in SSRT, and proportion of trigger failures) and metrics of the burst, we fit a generalized linear model using the MATLAB statistical toolbox, using the burst metric as a predictor.

We also consider bursts more broadly across a wider range of epochs. This approach compared the proportion of β-bursts that occurred during various epochs between canceled, non-canceled, and no-stop trials. These epochs included: fixation (−400 to −200 ms, pre-target), early post-action (100 to 300 ms, post-SSRT on canceled trials, or 100 to 300 ms, post saccade on no-stop and non-canceled trials), late post-action (−300 to −100 ms, pre-tone on canceled trials, or 400 to 600 ms, post-saccade on no-stop and non-canceled trials), and post-tone (100 to 300 ms, post-tone on all trial types). The p(bursts) was estimated at the individual contact level (n = 509). We compared activity between trial type and epoch using a two-way repeated-measures ANOVA.

### Neurometric approach

For the neurometric analysis, we calculated the proportion of bursts in the last 50 ms of the stop period on canceled trials (−50 to 0 ms, relative to SSRT). For each SSD, we subtracted the proportion of trials with a β-burst (neurometric) from the probability of canceling a saccade (psychometric measure). For the shuffled condition, we subtracted the proportion of trials with a β-burst at another SSD other than the one of interest. We then used these differences between the psychometric and neurometric values to calculate the sum of squared differences for both the shuffled and observed conditions. These values were then compared using an independent groups t-test, adjusting the p-value for multiple comparisons. We repeated this from a 1x to 10 x threshold of the median baseline power for the given session.

### Relating β-bursts to proactive control

To probe how baseline p(bursts) varied with the outcome of the previous trial, we compared p(β-bursts) occurring in the same fixation period (−400 to −200 ms) in no-stop trials following no-stop, canceled, and non-canceled trials. We compared activity between preceding trial types using a one-way repeated-measures ANOVA.

Finally, we look to see if β-bursts may influence the RT on a given trial. We took trials in which a movement was generated (correctly or incorrectly; no-stop and non-canceled trials, respectively), and then vincentized these trials into six quantiles (i.e., from fast RTs to slow RTs, binned in 20% bins from 0 to 100% of RTs). We then calculated the p(bursts) that occurred within the baseline (−400 to −200 ms, target) and immediately before the saccade (−200 to 0 ms, saccade) of the trials within each quantile. We then fit a generalized linear model using the MATLAB statistical toolbox, using the vincentized response time as a predictor.

### Comparing β-bursts during errors

We compared the proportion of bursts between trials in which a movement was correctly generated (no-stop) and those in which one was incorrectly generated (non-canceled error). We did this for two separate periods following the saccade: an early period (100 to 300 ms, post-saccade), and a late period (400 to 600 ms, post-saccade), for each contact within the cortex (n = 509). We compared activity between trial type and early/late error periods using a two-way repeated-measures ANOVA.

### Relating error-related β-burst activity to RT adaptation

We first identified no-stop trials following errors. We then divided these trials into two groups: ones in which the error trial had a burst within the 400 to 600 ms post-error period, and those that didn’t. We then identified the degree of post-error slowing by subtracting the RT on the no-stop trial from that on the proceeding non-canceled trial. We then compared whether this change in RT was significantly different between those trials in which a burst occurred, and those that didn’t, using an independent groups t-test. For this analysis, the alpha value was set at 0.05/509 (the number of contacts) to account for multiple comparisons. Similar results were observed at an alpha level of 0.05.

### Comparing β-power across layers

We computed the power spectral density to determine the relative power of the β oscillatory activity across cortical depth (Bastos et al., 2018; Maier et al., 2010; Westerberg et al., 2019). LFP was filtered between 1 and 200 Hz with a 4^th^-order bidirectional Butterworth filter and full-wave rectified to estimate power at each frequency. Power was normalized relative to the mean of a given column’s power at each frequency. We also computed the average β-power during the early post-action period (100 to 300 ms, post-SSRT) for each cortical contact within our laminar penetrations (n = 249 contacts), using the bandpower function of the MATLAB signal processing toolbox. For each session, we then normalized the power by denoting the power for each contact(/depth) as a function of the maximal β-power in the penetration. We then averaged power across all contacts within a given cortical layer. This normalized activity was then compared across sessions, and trial types. We compared activity between trial type and cortical layer using a two-way mixed-measures ANOVA, with layer as an independent factor, and trial type as a repeated factor.

### Comparing β-bursts across layers

We computed the p(bursts) observed in canceled trials across several epochs for each cortical contact within our laminar penetrations (n = 249 contacts). We focused on four epochs: fixation (−400 to −200 ms, pre-target), early post-action (100 to 300 ms, post-SSRT), late post-action (−300 to −100 ms, pre-tone), and post-tone (100 to 300 ms, post-tone). We then averaged p(bursts) across all contacts within a given cortical layer. We used a two-way mixed-measures ANOVA to compare activity across sessions, with layer as an independent factor, and trial type as a repeated factor.

We then looked at the spatiotemporal pattern of β-bursts by considering the BBDF across layers and times. We averaged the BBDF on canceled trials for each cortical contact within our laminar penetrations (n = 249 contacts), for each epoch of interest. Each contact was assigned to the corresponding cortical depth at which it was recorded. We then averaged this depth-aligned BBDF across sessions. Finally, we then normalized the time-depth BBDF by calculating the Z-score value for each depth-time point, relative to the mean and SD of the entire array for the epoch.

### Linking intracranial β-bursts to EEG β-bursts

We first looked at the p(burst) varied across epochs, dependent on the spatial scale of measuring bursts. We considered bursts in the EEG signal and three levels of the intracortical signal: the probability of observing a burst on individual intracortical contact, the probability of observing a burst on any of the intracortical contacts in each laminar layer (upper or lower), and the probability of observing a burst on any of the intracortical channels. We focused on six epochs: fixation (−400 to −200 ms, pre-target, canceled), target (0 to 200 ms, post-target, canceled), stopping (0 to 200 ms, post stop-signal-delay, canceled), early post-action (100 to 300 ms, post-saccade, non-canceled), late post-action (400 to 600 ms, post-saccade, non-canceled) and post-tone (100 to 300 ms, post-tone).

We then found trials in which bursts occurred in both the EEG and cortex, for each contact in our perpendicular penetrations (n = 249 contacts). Once we identified these trials, we determined the temporal difference between the time of the burst peak in the cortex, relative to the time of the EEG burst. We generated a shuffle condition by randomizing the trials from which the LFP bursts were sampled. Negative values represent bursts in the cortex prior to the EEG burst, and positive values represent bursts in the cortex following the EEG burst. These latency differences were then into 10 ms bins between −250 to 250 ms, across contacts in the upper and lower cortical layers. We then subtracted the shuffled from the observed p(bursts) in each bin.

To probe whether bursts in the cortex led those in the EEG, we calculated the area under the curve by multiplying the bin width (10 ms) by the observed-shuffled p(burst | bin). We then summed bins in the −50 to 0 ms, and 0 to 50 ms period relative to the EEG burst. We whether this area was greater than zero using a one-sample t-test, for both the upper and lower layers.

We then calculated the odds ratio to quantify the relationship between the observation of β-bursts in the EEG and LFP signals. LFP signals were considered at the individual channel level. To achieve this, we counted the number of trials in which bursts were observed in each window for (a) both the LFP and EEG signal, (b) neither the LFP nor EEG signal, (c) just the LFP signal, and (d) just the EEG signal. Based on our analysis, we used 50 ms bins. These bins ranged from −600 to 200 ms relative to the target onset on canceled trials, 0 to 800 ms following the saccade on non-canceled trials, −200 to 600 ms relative to the stop-signal on canceled trials, and −600 to 200 ms relative to the feedback tone on canceled trials. We then conducted a Fisher’s exact test to determine whether the bursts were significantly co-incidental. To achieve this, we conducted a right-tailed test reflecting an alternative hypothesis that the odds ratio is greater than 1; this would represent a greater co-incidence of a burst in cortex and EEG. This analysis used an alpha level set at 0.003 (0.05/17 bins). We present results as an average of the difference between observed and shuffled bursts, across all contacts in our perpendicular penetration (n = 249 contacts).

Finally, we repeated this analysis by comparing the intracortical signals. Across the same windows and epochs as above, we counted the number of trials in which bursts were observed in each window for (a) both channels, (b) neither channel, (c) just channel one, and (d) just channel two. We then align the relevant channels to their respective depth in the cortex and, for each epoch, average this activity across all time bins and all sessions. To determine intralaminar (i.e., within a layer) and interlaminar (i.e. between layers) differences, we averaged the odds ratios within these conditions. We then compared these odds ratios between conditions using a two-way independent-group ANOVA (epoch and laminar-congruence).

### Data and Software Availability

MATLAB code for the analyses conducted in this study is openly available on GitHub (https://github.com/stevenerrington/2021-sef-cortical-beta) and processed data is available through OSF (https://osf.io/5rnk6/). Requests for materials should be addressed to S.E.P. or J.D.S.

## Acknowledgments

This work was supported by National Institutes of Health Grants R01-MH-55806, R01-EY-019882, P30-EY-008126, P30-HD-01505, F31-EY-031293, T32-EY-007135, NSERC RGPIN-2022-04592, and Robin and Richard Patton through the E. Bronson Ingram Chair in Neuroscience. We thank B. Williams, R. Williams, M. Maddox, M.S. Schall, I. Haniff, S. Motorny, D. Richardson, L. Toy, and M.R. Feurtado, for technical support. We also thank G. Logan, K.A. Lowe, U. Rutishauser, & T. Womelsdorf, for useful conversations regarding the work.

## Author Contributions

J.D.S. and G.F.W. oversaw data collection. S.P.E. conceived and conducted the analyses. J.A.W. contributed resources for laminar analyses. S.P.E., J.A.W., G.F.W., and J.D.S. wrote the paper. All authors approved the final version of this report.

## Declaration of interests

The authors declare no competing interests

## Notes

### Competing Interest Statement

The authors have declared no competing interest.

## REFERENCES

Bastos, A.M., Loonis, R., Kornblith, S., Lundqvist, M., and Miller, E.K. (2018). Laminar recordings in frontal cortex suggest distinct layers for maintenance and control of working memory. Proc Natl Acad Sci U S A 115, 1117–1122.

Bimbi, M., Festante, F., Coude, G., Vanderwert, R.E., Fox, N.A., and Ferrari, P.F. (2018). Simultaneous scalp recorded EEG and local field potentials from monkey ventral premotor cortex during action observation and execution reveals the contribution of mirror and motor neurons to the mu-rhythm. Neuroimage 175, 22–31.

Bonini, F., Burle, B., Liegeois-Chauvel, C., Regis, J., Chauvel, P., and Vidal, F. (2014). Action monitoring and medial frontal cortex: leading role of Supplementary motor area. Science 343, 888–891.

Boucher, L., Palmeri, T.J., Logan, G.D., and Schall, J.D. (2007a). Inhibitory control in mind and brain: an interactive race model of countermanding saccades. Psychol Rev 114, 376–397.

Boucher, L., Stuphorn, V., Logan, G.D., Schall, J.D., and Palmeri, T.J. (2007b). Stopping eye and hand movements: are the processes independent? Percept Psychophys 69, 785–801.

Brown, J.W., Hanes, D.P., Schall, J.D., and Stuphorn, V. (2008). Relation of frontal eye field activity to saccade initiation during a countermanding task. Exp Brain Res 190, 135–151.

Cabel, D.W., Armstrong, I.T., Reingold, E., and Munoz, D.P. (2000). Control of saccade initiation in a countermanding task using visual and auditory stop signals. Exp Brain Res 133, 431–441.

Castiglione, A., Wagner, J., Anderson, M., and Aron, A.R. (2019). Preventing a Thought from Coming to Mind Elicits Increased Right Frontal Beta Just as Stopping Action Does. Cereb Cortex 29, 2160–2172.

Cohen, J.Y., Heitz, R.P., Schall, J.D., and Woodman, G.F. (2009). On the origin of event-related potentials indexing covert attentional selection during visual search. J Neurophysiol 102, 2375–2386.

Cohen, M.X. (2014). A neural microcircuit for cognitive conflict detection and signaling. Trends Neurosci 37, 480–490.

Colonius, H., Ozyurt, J., and Arndt, P.A. (2001). Countermanding saccades with auditory stop signals: testing the race model. Vision Res 41, 1951–1968.

Cosman, J.D., Lowe, K.A., Zinke, W., Woodman, G.F., and Schall, J.D. (2018). Prefrontal Control of Visual Distraction. Curr Biol 28, 1330.

Diesburg, D.A., Greenlee, J.D., and Wessel, J.R. (2021). Cortico-subcortical beta burst dynamics underlying movement cancellation in humans. Elife 10.

Diesburg, D.A., and Wessel, J.R. (2021). The Pause-then-Cancel model of human action-stopping: Theoretical considerations and empirical evidence. Neurosci Biobehav Rev 129, 17–34.

Emeric, E.E., Leslie, M., Pouget, P., and Schall, J.D. (2010). Performance monitoring local field potentials in the medial frontal cortex of primates: Supplementary eye field. J Neurophysiol 104, 1523–1537.

Enz, N., Ruddy, K.L., Rueda-Delgado, L.M., and Whelan, R. (2021). Volume of beta-Bursts, But Not Their Rate, Predicts Successful Response Inhibition. J Neurosci 41, 5069–5079.

Errington, S.P., Woodman, G.F., and Schall, J.D. (2020). Dissociation of Medial Frontal beta-Bursts and Executive Control. J Neurosci 40, 9272–9282.

Fujii, N., Mushiake, H., and Tanji, J. (2002). Distribution of eye- and arm-movement-related neuronal activity in the SEF and in the SMA and Pre-SMA of monkeys. J Neurophysiol 87, 2158–2166.

Godlove, D.C., Garr, A.K., Woodman, G.F., and Schall, J.D. (2011). Measurement of the extraocular spike potential during saccade countermanding. J Neurophysiol 106, 104–114.

Godlove, D.C., Maier, A., Woodman, G.F., and Schall, J.D. (2014). Microcircuitry of agranular frontal cortex: Testing the generality of the canonical cortical microcircuit. J Neurosci 34, 5355–5369.

Godlove, D.C., and Schall, J.D. (2016). Microsaccade production during saccade cancelation in a stop-signal task. Vision Res 118, 5–16.

Hanes, D.P., and Carpenter, R.H. (1999). Countermanding saccades in humans. Vision Res 39, 2777–2791.

Hanes, D.P., and Schall, J.D. (1995). Countermanding saccades in macaque. Vis Neurosci 12, 929–937.

Hannah, R., Muralidharan, V., Sundby, K.K., and Aron, A.R. (2020). Temporally-precise disruption of prefrontal cortex informed by the timing of beta bursts impairs human action-stopping. Neuroimage 222, 117222.

Hanslmayr, S., Matuschek, J., and Fellner, M.C. (2014). Entrainment of prefrontal beta oscillations induces an endogenous echo and impairs memory formation. Curr Biol 24, 904–909.

Heitz, R.P., Cohen, J.Y., Woodman, G.F., and Schall, J.D. (2010). Neural correlates of correct and errant attentional selection revealed through N2pc and frontal eye field activity. J Neurophysiol 104, 2433–2441.

Herrera, B., Sajad, A., Woodman, G.F., Schall, J.D., and Riera, J.J. (2020). A Minimal Biophysical Model of Neocortical Pyramidal Cells: Implications for Frontal Cortex Microcircuitry and Field Potential Generation. J Neurosci 40, 8513–8529.

Herrera, B., Westerberg, J.A., Schall, M.S., Maier, A., Woodman, G.F., Schall, J.D., and Riera, J.J. (2022). Resolving the mesoscopic missing link: Biophysical modeling of EEG from cortical columns in primates. Neuroimage 263, 119593.

Jana, S., Hannah, R., Muralidharan, V., and Aron, A.R. (2020). Temporal cascade of frontal, motor and muscle processes underlying human action-stopping. Elife 9.

Kawagoe, R., Takikawa, Y., and Hikosaka, O. (1998). Expectation of reward modulates cognitive signals in the basal ganglia. Nat Neurosci 1, 411–416.

Kornylo, K., Dill, N., Saenz, M., and Krauzlis, R.J. (2003). Cancelling of pursuit and saccadic eye movements in humans and monkeys. J Neurophysiol 89, 2984–2999.

Lo, C.C., Boucher, L., Pare, M., Schall, J.D., and Wang, X.J. (2009). Proactive inhibitory control and attractor dynamics in countermanding action: a spiking neural circuit model. J Neurosci 29, 9059–9071.

Logan, G.D., and Cowan, W.B. (1984). On the ability to inhibit thought and action - a theory of an act of control. Psychol Rev 91, 295–327.

Logan, G.D., and Irwin, D.E. (2000). Don’t look! Don’t touch! Inhibitory control of eye and hand movements. Psychon Bull Rev 7, 107–112.

Logan, G.D., Yamaguchi, M., Schall, J.D., and Palmeri, T.J. (2015). Inhibitory control in mind and brain 2.0: blocked-input models of saccadic countermanding. Psychol Rev 122, 115–147.

Lundqvist, M., Rose, J., Herman, P., Brincat, S.L., Buschman, T.J., and Miller, E.K. (2016). Gamma and Beta Bursts Underlie Working Memory. Neuron 90, 152–164.

Maier, A., Adams, G.K., Aura, C., and Leopold, D.A. (2010). Distinct superficial and deep laminar domains of activity in the visual cortex during rest and stimulation. Front Syst Neurosci 4.

Martinez-Trujillo, J.C., Medendorp, W.P., Wang, H., and Crawford, J.D. (2004). Frames of reference for eye-head gaze commands in primate Supplementary eye fields. Neuron 44, 1057–1066.

Matzke, D., Dolan, C.V., Logan, G.D., Brown, S.D., and Wagenmakers, E.J. (2013a). Bayesian parametric estimation of stop-signal reaction time distributions. J Exp Psychol Gen 142, 1047–1073.

Matzke, D., Love, J., and Heathcote, A. (2017). A bayesian approach for estimating the probability of trigger failures in the stop-signal paradigm. Behav Res Methods 49, 267–281.

Matzke, D., Love, J., Wiecki, T.V., Brown, S.D., Logan, G.D., and Wagenmakers, E.J. (2013b). Release the BEESTS: Bayesian Estimation of Ex-Gaussian STop-Signal reaction time distributions. Front Psychol 4, 918.

Morein-Zamir, S., and Kingstone, A. (2006). Fixation offset and stop signal intensity effects on saccadic countermanding: a crossmodal investigation. Exp Brain Res 175, 453–462.

Mosher, C.P., Mamelak, A.N., Malekmohammadi, M., Pouratian, N., and Rutishauser, U. (2021). Distinct roles of dorsal and ventral subthalamic neurons in action selection and cancellation. Neuron 109, 869–881 e866.

Muralidharan, V., Aron, A.R., and Schmidt, R. (2022). Transient beta modulates decision thresholds during human action-stopping. Neuroimage 254, 119145.

Naess, S., Halnes, G., Hagen, E., Hagler, D.J., Jr., Dale, A.M., Einevoll, G.T., and Ness, T.V. (2021). Biophysically detailed forward modeling of the neural origin of EEG and MEG signals. Neuroimage 225, 117467.

Ninomiya, T., Dougherty, K., Godlove, D.C., Schall, J.D., and Maier, A. (2015). Microcircuitry of agranular frontal cortex: contrasting laminar connectivity between occipital and frontal areas. J Neurophysiol 113, 3242–3255.

Passingham, R.E., Bengtsson, S.L., and Lau, H.C. (2010). Medial frontal cortex: from self-generated action to reflection on one’s own performance. Trends Cogn Sci 14, 16–21.

Purcell, B.A., Schall, J.D., and Woodman, G.F. (2013). On the origin of event-related potentials indexing covert attentional selection during visual search: timing of selection by macaque frontal eye field and event-related potentials during pop-out search. J Neurophysiol 109, 557–569.

Purcell, B.A., Weigand, P.K., and Schall, J.D. (2012). Supplementary eye field during visual search: salience, cognitive control, and performance monitoring. J Neurosci 32, 10273–10285.

Rae, C.L., Hughes, L.E., Weaver, C., Anderson, M.C., and Rowe, J.B. (2014). Selection and stopping in voluntary action: a meta-analysis and combined fMRI study. Neuroimage 86, 381–391.

Rushworth, M.F., Walton, M.E., Kennerley, S.W., and Bannerman, D.M. (2004). Action sets and decisions in the medial frontal cortex. Trends Cogn Sci 8, 410–417.

Sajad, A., Errington, S.P., and Schall, J.D. (In Press). Functional architecture of executive control and associated event-related potentials in macaques. Nat Commun.

Sajad, A., Godlove, D.C., and Schall, J.D. (2019). Cortical microcircuitry of performance monitoring. Nat Neurosci 22, 265–274.

Sandhaeger, F., von Nicolai, C., Miller, E.K., and Siegel, M. (2019). Monkey EEG links neuronal color and motion information across species and scales. Elife 8.

Scangos, K.W., and Stuphorn, V. (2010). Medial frontal cortex motivates but does not control movement initiation in the countermanding task. J Neurosci 30, 1968–1982.

Schall, J.D. (1991). Neuronal activity related to visually guided saccades in the frontal eye fields of rhesus monkeys: comparison with Supplementary eye fields. J Neurophysiol 66, 559–579.

Schall, J.D., Stuphorn, V., and Brown, J.W. (2002). Monitoring and control of action by the frontal lobes. Neuron 36, 309–322.

Sharp, D.J., Bonnelle, V., De Boissezon, X., Beckmann, C.F., James, S.G., Patel, M.C., and Mehta, M.A. (2010). Distinct frontal systems for response inhibition, attentional capture, and error processing. Proc Natl Acad Sci U S A 107, 6106–6111.

Sherman, M.A., Lee, S., Law, R., Haegens, S., Thorn, C.A., Hamalainen, M.S., Moore, C.I., and Jones, S.R. (2016). Neural mechanisms of transient neocortical beta rhythms: Converging evidence from humans, computational modeling, monkeys, and mice. Proc Natl Acad Sci U S A 113, E4885–4894.

Shin, H., Law, R., Tsutsui, S., Moore, C.I., and Jones, S.R. (2017). The rate of transient beta frequency events predicts behavior across tasks and species. Elife 6.

Smith, M.A., and Sommer, M.A. (2013). Spatial and temporal scales of neuronal correlation in visual area V4. J Neurosci 33, 5422–5432.

Stuphorn, V., Brown, J.W., and Schall, J.D. (2010). Role of Supplementary eye field in saccade initiation: executive, not direct, control. J Neurophysiol 103, 801–816.

Stuphorn, V., Taylor, T.L., and Schall, J.D. (2000). Performance monitoring by the Supplementary eye field. Nature 408, 857–860.

Thakkar, K.N., Schall, J.D., Boucher, L., Logan, G.D., and Park, S. (2011). Response inhibition and response monitoring in a saccadic countermanding task in schizophrenia. Biol Psychiatry 69, 55–62.

Thakkar, K.N., Schall, J.D., Logan, G.D., and Park, S. (2015). Cognitive control of gaze in bipolar disorder and schizophrenia. Psychiatry Res 225, 254–262.

Thompson, K.G., Hanes, D.P., Bichot, N.P., and Schall, J.D. (1996). Perceptual and motor processing stages identified in the activity of macaque frontal eye field neurons during visual search. J Neurophysiol 76, 4040–4055.

Verbruggen, F., Aron, A.R., Band, G.P., Beste, C., Bissett, P.G., Brockett, A.T., Brown, J.W., Chamberlain, S.R., Chambers, C.D., Colonius, H., et al. (2019). A consensus guide to capturing the ability to inhibit actions and impulsive behaviors in the stop-signal task. Elife 8.

Walton, M.M., and Gandhi, N.J. (2006). Behavioral evaluation of movement cancellation. J Neurophysiol 96, 2011–2024.

Wattiez, N., Poitou, T., Rivaud-Pechoux, S., and Pouget, P. (2016). Evidence for spatial tuning of movement inhibition. Exp Brain Res 234, 1957–1966.

Wessel, J.R. (2020). beta-Bursts Reveal the Trial-to-Trial Dynamics of Movement Initiation and Cancellation. J Neurosci 40, 411–423.

Westerberg, J.A., Cox, M.A., Dougherty, K., and Maier, A. (2019). V1 microcircuit dynamics: altered signal propagation suggests intracortical origins for adaptation in response to visual repetition. J Neurophysiol 121, 1938–1952.

Westerberg, J.A., Maier, A., Woodman, G.F., and Schall, J.D. (2020). Performance Monitoring during Visual Priming. J Cogn Neurosci 32, 515–526.

Westerberg, J.A., Schall, M.S., Maier, A., Woodman, G.F., and Schall, J.D. (2022). Laminar microcircuitry of visual cortex producing attention-associated electric fields. Elife 11.

Whittingstall, K., and Logothetis, N.K. (2009). Frequency-band coupling in surface EEG reflects spiking activity in monkey visual cortex. Neuron 64, 281–289.

Wiecki, T.V., and Frank, M.J. (2013). A computational model of inhibitory control in frontal cortex and basal ganglia. Psychol Rev 120, 329–355.

Xing, D., Yeh, C.I., Burns, S., and Shapley, R.M. (2012). Laminar analysis of visually evoked activity in the primary visual cortex. Proc Natl Acad Sci U S A 109, 13871–13876.

Yu, Y., Escobar Sanabria, D., Wang, J., Hendrix, C.M., Zhang, J., Nebeck, S.D., Amundson, A.M., Busby, Z.B., Bauer, D.L., Johnson, M.D., et al. (2021). Parkinsonism Alters Beta Burst Dynamics across the Basal Ganglia-Motor Cortical Network. J Neurosci 41, 2274–2286.

