## Supplementary Information for "Cortical Contributions to Medial Frontal β-Bursts during Executive Control"

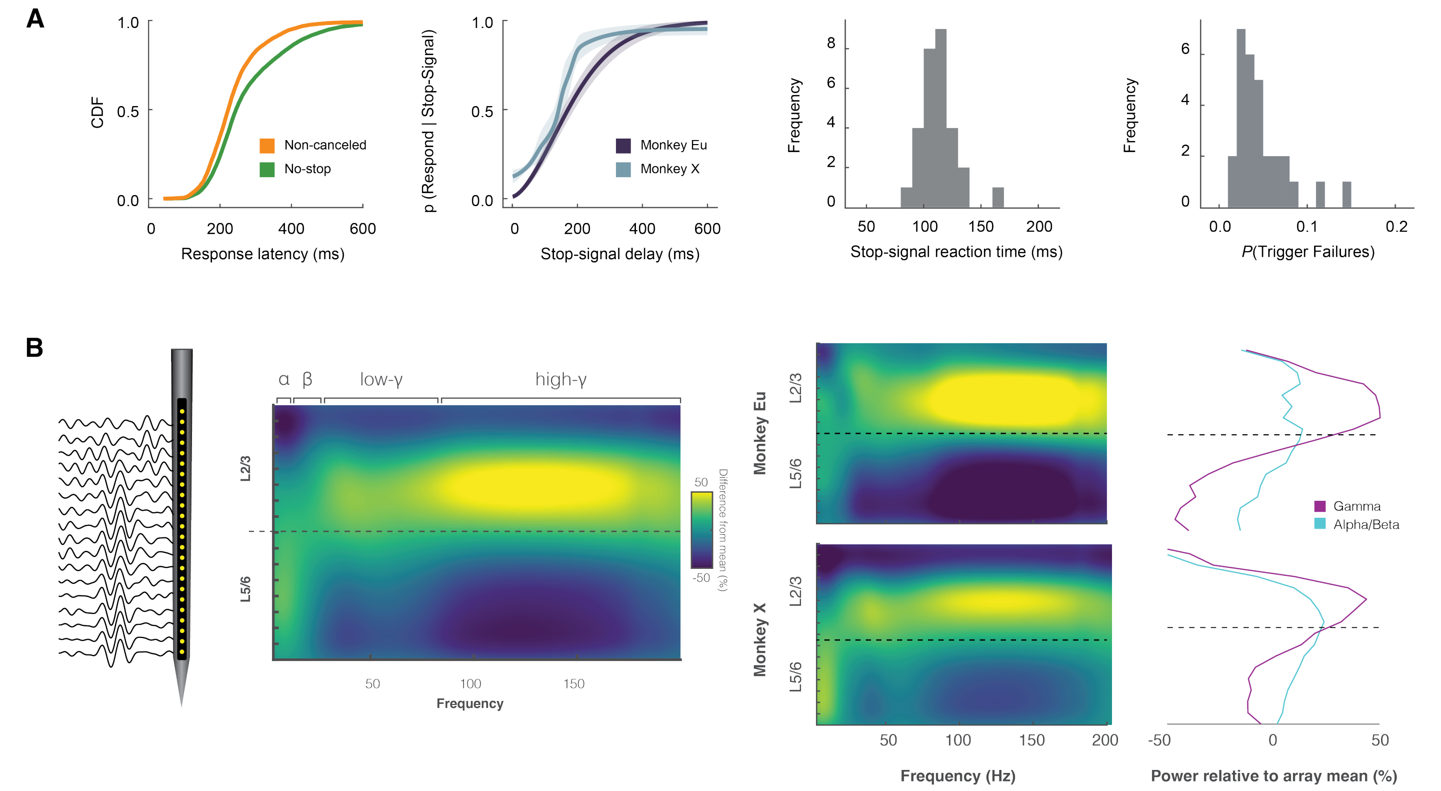


**Supplementary Fig. 1 | Countermanding behavior & experimental procedures.**

**A.** Saccade-countermanding behavior. *Left*, cumulative distribution function of response latencies on no-stop (green) and non-canceled (yellow) trials. Response latencies on non-canceled trials were faster than those on no-stop trials. *Center-left*: inhibition function plotting the probability of responding across stop-signal delays. Weibull functions were fitted to data from each session. The mean of these Weibull functions across sessions and the corresponding 95% CI is plotted for each monkey (monkey Eu: purple; X: blue). Errors increased with stop-signal delay. *Center-right*: distribution of mean stop-signal reaction times across sessions. Monkeys exhibited normal stop-signal reaction times (SSRT). *Right*: distribution of the proportion of trigger failures across sessions. Monkeys exhibited a low proportion of trigger failures.

**B**. Power spectral density. We computed the power spectral density to determine relative power of oscillatory activity across cortical depth. We observed high-frequencies (i.e. gamma) were most pronounced in upper cortical layers, whilst lower-frequencies (i.e. alpha) were most pronounced in lower cortical layers. β-activity was most powerful in the middle-to-lower cortical layers, around the L3/5 divide (dashed line) in SEF. This was observed across both monkeys.

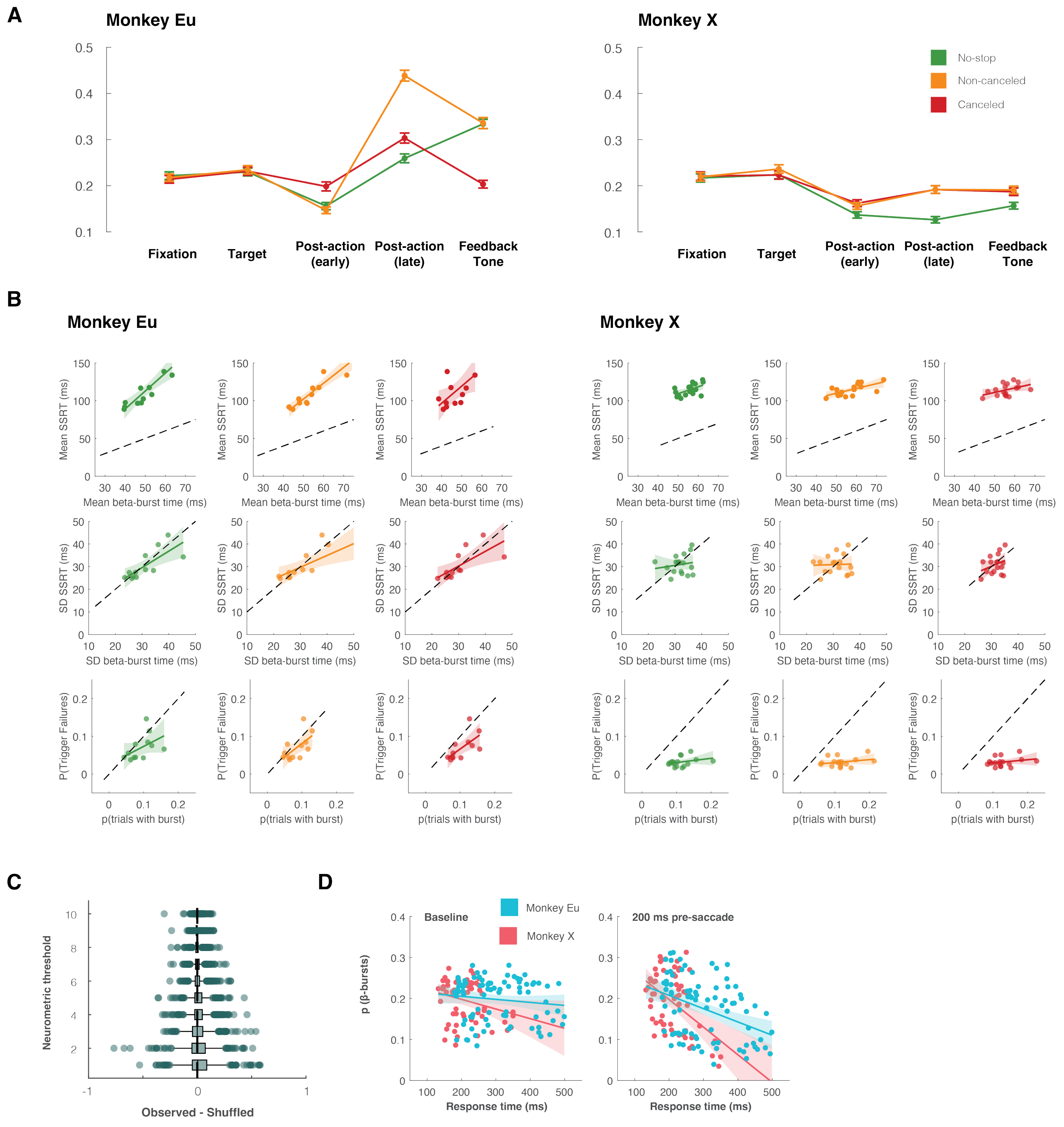


**Supplementary Fig. 2 | Functional properties of intracranial β-bursts.**

**A**. Mean ± SEM of probability of LFP β-bursts across different epochs on canceled (red), non-canceled (yellow) and no-stop (green) trials. LFP β-bursts varied across epochs. Bursts were most common following actions for both monkeys.

**B.** Association between properties of a burst during the stopping period (i.e. onset, variation, p(burst)) against metrics of stopping behavior (i.e. mean SSRT, SD SSRT, and P(trigger failures)), for canceled, no-stop, and non-canceled trials. Regression lines are presented with a 95% CI for each correlation.

**C**. Difference between the sum of squared differences for the observed and shuffled conditions, for each neurometric threshold. We varied the median detection threshold for defining a burst between one and ten times the median power of all contacts in a session. In no sessions were LFP β-bursts observed commonly enough to account for response inhibition.

**D**. Relationship between mean p(β-burst) and means of session-wise response time quantiles collapsed across all sessions. Bursts were calculated in a baseline period (-400 to -200 ms, pre-target; left) and a 200 ms preceding saccade initiation (right). Regression line with CI demonstrates significant negative association during the pre-saccade, but not the baseline period, for both monkeys.

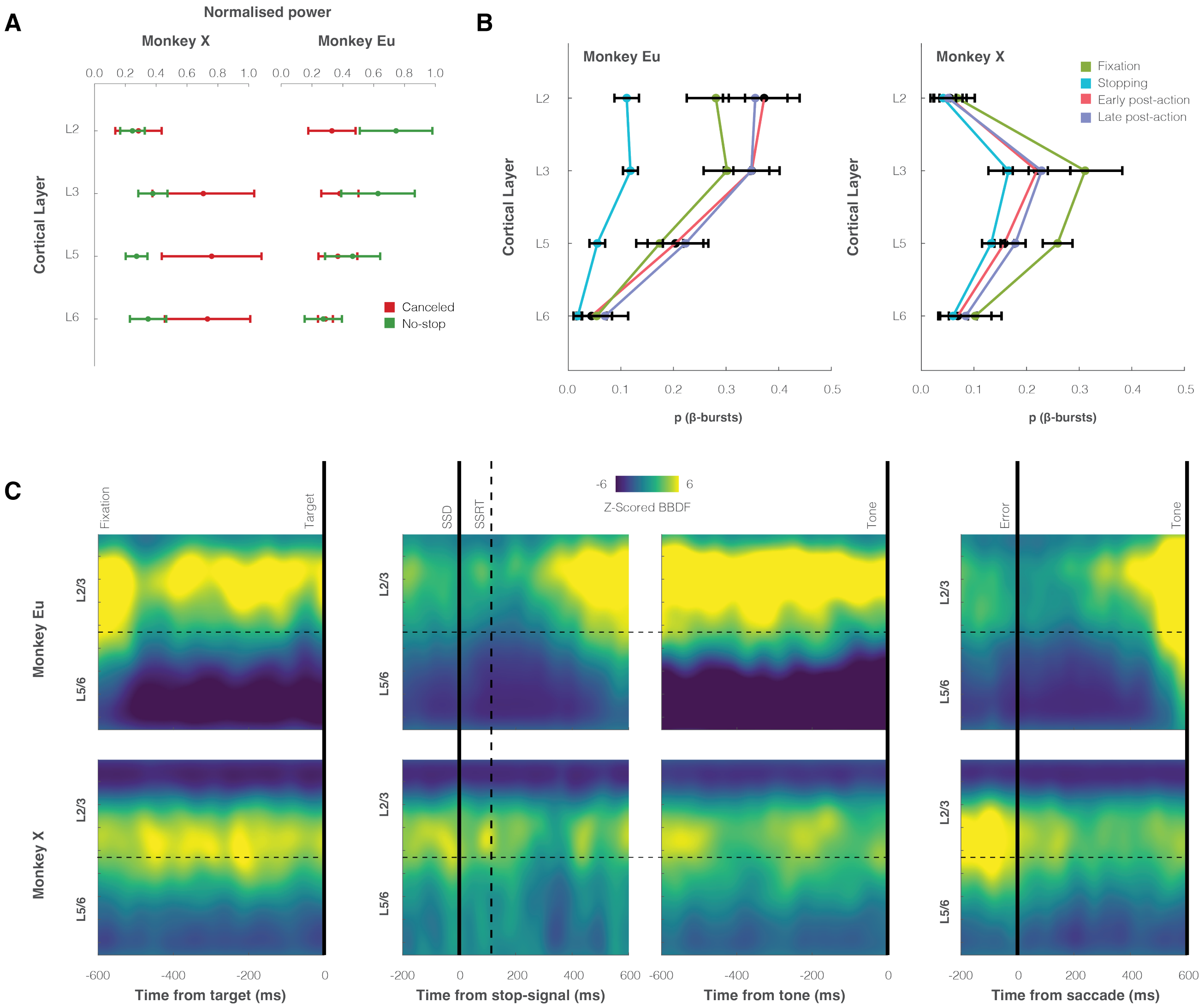


**Supplementary Fig. 3 | Laminar properties of intracranial β-bursts.**

**A**. Normalized power in each cortical layer during response inhibition for canceled and latency-matched no-stop trials. The elevation of LFP β power during response inhibition happened in the deep layers. Conventions as in Fig. 1.

**B**. Proportion of (β-bursts) in canceled trials across four trial epochs (fixation, green; stopping, blue; early post-action, pink; late post-action, purple) and cortical layers. Variability was observed across two monkeys, but bursts were most common during all epochs in L3 for both monkeys.

**C**. Heat maps of LFP β-burst incidence across time and cortical layers on canceled trials during foreperiod (left), stopping and early post-action period (left-center), late post-action period (right-center), and error (right) epochs. Yellow represents a greater proportion of (β-bursts) at a given time and depth, relative to other times and depths.

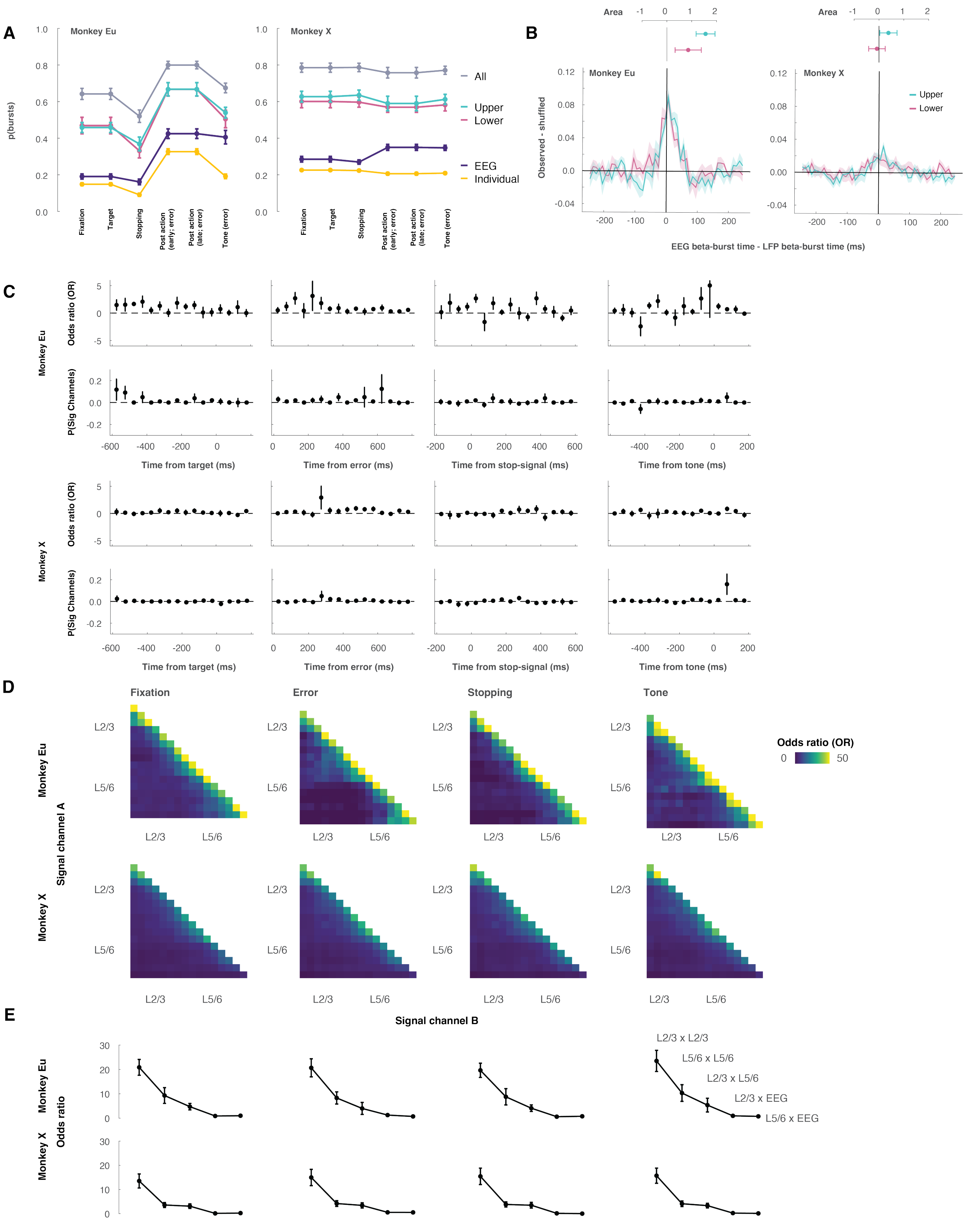


### Supplementary Fig. 4 | Comparing EEG and LFP β-bursts.

**A**. Mean ± SEM of p(β-burst) counted within epochs in individual LFP (gold) and EEG (purple) contacts and summed across contacts in L2/3 (teal), in L5/6 (magenta), and from L2 to L6 (grey). The number of LFP β-bursts within a cortical column exceeds the number of EEG β-bursts.

**B**. Distribution after shuffle subtraction of shortest times between EEG β-bursts and LFP β-bursts detected in L2/3 (teal) and in L5/6 (magenta) plotted with positive (negative) values for LFP β-bursts following (preceding) EEG β-bursts. Only ~5% of EEG β-bursts were associated more than chance with LFP β-bursts, and this association was observed only within ±50 ms of the EEG β-burst. Mean ± SEM area under the distributions across sessions, plotted above, indicates a weak tendency for LFP β-bursts in L2/3 but not L5/6 to follow EEG β-bursts.

**C**. Mean ± SEM of shuffle-corrected odds ratio of detecting an LFP β-burst ±50 ms from occurrence of an EEG β-burst (above) and of probability of detected significant odds ratio > 1 (below) during response inhibition. Coincident presence or absence of β-bursts across EEG and LFP channels was vanishingly rare.

**D**. Color map of odds ratios of LFP and EEG β-bursts shows association in adjacent but not distant contacts.

**E**. Mean ± SEM odds ratios of coincidences of β-bursts occurring within L2/3 LFP channels, within L5/6 LFP channels, across L2/3 and L5/6 LFP, and across EEG and LFP in L2/3 and in L5/6. Odds ratio scaled with prevalence and was effectively nil across EEG and LFP channels.

**Supplementary Table 1.** $R^2$ (significance values) between beta-burst and behavioral stopping metrics, across trial types.

| | | **Trial Type** | | |
|---|---|---|---|---|
| **Monkey** | **SSRT Metric** | **No-stop** | **Non-canceled** | **Canceled** |
| | Mean | **0.542 (p < 0.001)** | **0.630 (p < 0.001)** | 0.197 (p = 0.016) |
| All | SD | 0.299 (p = 0.002) | 0.237 (p = 0.007) | **0.339 (p < 0.001)** |
| | Trigger Failure | 0.007 (p = 0.674) | 0.001 (p = 0.888) | 0.011 (p = 0.593) |
| | Mean | **0.760 (p < 0.001)** | **0.865 (p < 0.001)** | 0.393 (p = 0.029) |
| Eu | SD | 0.575 (p = 0.004) | 0.517 (p = 0.008) | 0.497 (p = 0.011) |
| | Trigger Failure | 0.226 (p = 0.118) | 0.374 (p = 0.035) | 0.425 (p = 0.022) |
| | Mean | 0.308 (p = 0.021) | 0.416 (p = 0.005) | 0.206 (p = 0.068) |
| X | SD | 0.028 (p = 0.523) | 0.001 (p = 0.886) | 0.103 (p = 0.210) |
| | Trigger Failure | 0.087 (p = 0.251) | 0.070 (p = 0.303) | 0.067 (p = 0.315) |

**Bold values** signify $p < 0.001$ (Bonferroni adjusted)

Mean represents mean maximal burst time x mean SSRT; SD represents SD maximal burst time x SD SSRT; Trigger Failure represents p(bursts) x p(TF).

**Supplementary Table 2.** Metrics (Mean ± SEM) of beta-burst dynamics during the STOP period across all contacts (n = 509)

| | | **Trial Type**<br>**Canceled** | **No-stop** | **Non-canceled** |
|---|---|---|---|---|
| All<br>(n = 509) | P(Burst) (%) | 12.2 ± 0.4 | 10.6 ± 0.4 | 10.8 ± 0.4 |
| | Frequency (Hz) | 19.8 ± 0.3 | 20.2 ± 0.3 | 19.4 ± 0.3 |
| | Peak time (ms) | 52.2 ± 0.5 | 53.4 ± 0.4 | 56.4 ± 0.6 |
| | Onset (ms) | -179.3 ± 2.5 | -175.7 ± 2.7 | -175. 6 ± 2.7 |
| | Offset (ms) | 173.7 ± 2.4 | 165.9 ± 1.9 | 175.0 ± 2.7 |
| | Duration (ms) | 353.0 ± 4.5 | 341.6 ± 4.0 | 350.6 ± 4.5 |
| | Volume ($10^{11}$) | 2.448 ± 0.178 | 2.442 ± 0.278 | 2.615 ± 0.411 |
| Eu<br>(n = 217) | P(Burst) | 10.8 ± 0.5 | 9.5 ± 0.5 | 8.7 ± 0.5 |
| | Frequency (Hz) | 20.6 ± 0.5 | 21.8 ± 0.5 | 21.3 ± 0.5 |
| | Peak time (ms) | 46.9 ± 0.7 | 50.2 ± 0.7 | 55.3 ± 1.0 |
| | Onset (ms) | -152.7 ± 2.0 | -147.0 ± 1.7 | -146.1 ± 2.8 |
| | Offset (ms) | 149.5 ± 1.8 | 148.6 ± 2.5 | 154.5 ± 3.8 |
| | Duration (ms) | 302.2 ± 3.0 | 295.6 ± 3.4 | 300.6 ± 5.1 |
| | Volume ($10^{11}$) | 2.084 ± 0.052 | 2.032 ± 0.080 | 1.924 ± 0.101 |
| X<br>(n = 292) | P(Burst) | 13.3 ± 0.6 | 11.5 ± 0.5 | 12.3 ± 0.6 |
| | Frequency (Hz) | 19.2 ± 0.4 | 19.0 ± 0.4 | 17.9 ± 0.3 |
| | Peak time (ms) | 56.2 ± 0.6 | 55.7 ± 0.5 | 57.2 ± 0.7 |
| | Onset (ms) | -199.4 ± 3.7 | -197.1 ± 4.0 | -197.4 ± 3.8 |
| | Offset (ms) | 191.9 ± 3.6 | 178.7 ± 2.4 | 190.0 ± 3.5 |
| | Duration (ms) | 391.3 ± 6.8 | 375.8 ± 5.7 | 387.5 ± 6.1 |
| | Volume ($10^{11}$) | 2.722 ± 0.310 | 2.747 ± 0.481 | 3.125 ± 0.710 |
